# Experimental evolution-driven identification of Arabidopsis rhizosphere competence genes in *Pseudomonas protegens*

**DOI:** 10.1101/2020.12.01.407551

**Authors:** Erqin Li, Hao Zhang, Henan Jiang, Corné M.J. Pieterse, Alexandre Jousset, Peter A.H.M. Bakker, Ronnie de Jonge

## Abstract

Beneficial plant root-associated microorganisms carry out a range of functions that are essential for plant performance. Establishment of a bacterium on plant roots, however, requires overcoming several challenges, including competition with neighboring microorganisms and host immunity. Forward and reverse genetics has led to the identification of mechanisms that are used by beneficial microorganisms to overcome these challenges such as the production of iron-chelating compounds, the formation of strong biofilms, or the concealment of characteristic microbial molecular patterns that trigger the host immune system. However, how such mechanisms arose from an evolutionary perspective is much less understood. To study bacterial adaptation in the rhizosphere, we employed experimental evolution to track the physiological and genetic dynamics of root-dwelling *Pseudomonas protegens* in the *Arabidopsis thaliana* rhizosphere under axenic conditions. This simplified binary one plant-one bacterium system allows for the amplification of key adaptive mechanisms for bacterial rhizosphere colonization. We identified 35 mutations, including single-nucleotide polymorphisms, insertions, and deletions, distributed over 28 genes. We found that mutations in genes encoding global regulators, and in genes for siderophore production, cell surface decoration, attachment, and motility accumulated in parallel, underlining that bacterial adaptation to the rhizosphere follows multiple strategies. Notably, we observed that motility increased in parallel across multiple independent evolutionary lines. Altogether these results underscore the strength of experimental evolution to identify key genes, pathways, and processes for bacterial rhizosphere colonization, and a methodology for the development of elite beneficial microorganisms with enhanced root-colonizing capacities that can support sustainable agriculture in the future.

**Importance:** Beneficial root-associated microorganisms carry out many functions that are essential for plant performance. Establishment of a bacterium on plant roots, however, requires overcoming many challenges. Previously, diverse mechanisms that are used by beneficial microorganisms to overcome these challenges were identified. However, how such mechanisms have developed from an evolutionary perspective is much less understood. Here, we employed experimental evolution to track the evolutionary dynamics of a root-dwelling pseudomonad on the root of Arabidopsis. We find that mutations in global regulators, as well as in genes for siderophore production, cell surface decoration, attachment, and motility accumulate in parallel, underlining these strategies for bacterial adaptation to the rhizosphere. We identified 35 mutations distributed over 28 genes. Altogether our results demonstrate the power of experimental evolution to identify key pathways for rhizosphere colonization and a methodology for the development of elite beneficial microorganisms that can support sustainable agriculture.

## Introduction

Plants are associated with complex microbial communities assembled into a functional microbiome that safeguards optimal plant performance under harsh environmental conditions (1). The rhizosphere is a particularly interesting hotspot of plant-microbe interactions. Plants deposit up to 44% of their photosynthetically fixed carbon into the rhizosphere, fueling a specific microbial community (2). The microbial species pool in the bulk soil is the source from which the root microbiome is recruited, and plant genotype, immune responses, and environmental factors are postulated to affect this process (3–6). The establishment of beneficial microbial associations requires a high degree of coordination of both plant and microbial responses by means of a continuous molecular dialogue (7, 8). Plant-associated microorganisms can improve plant yield by protecting the plant from abiotic stresses (9), improving plant nutrition and growth (10–12), antagonizing soil-borne pathogens (13), or stimulating plant immunity (14). To exert their beneficial effects on plant performance, bacteria must colonize the roots efficiently and establish significant populations. For example, bacterial population densities above 10^5^ cells per gram of root are required for efficient suppression of soil-borne plant pathogens by *Pseudomonas* spp. (15, 16). Therefore, bacterial adaptation to the plant root environment may be essential for the successful implementation of microbiome services in agriculture in order to support plant health.

Among all root-dwelling organisms, fluorescent *Pseudomonas* spp. are well characterized in terms of the traits required for their growth in the rhizosphere and for the establishment of beneficial plant-microbe interactions (17, 18). To study bacterial traits involved in efficient root colonization, mutants defective in specific traits suspected to be involved in colonization, are compared to the parental strains for their ability to colonize plant roots. Such studies have highlighted a range of bacterial traits involved in efficient root colonization, including flagella (19), surface lipopolysaccharides (LPS) (20), and amino acid synthesis (21). Using random mutagenesis and by determining the fitness of each mutant in competition with its parental strain in the rhizosphere, many other important traits for rhizosphere competence in *Pseudomonas* were discovered (17). Recently, random mutagenesis in *Pseudomonas capeferrum* WCS358 led to the identification of two genes that are important for gluconic acid (GA) biosynthesis. GA, in turn, is essential for the suppression of local, flagellin-induced root immune responses (22). Such suppression was shown to be important for rhizosphere competence as GA-deficient mutants maintain reduced populations in the rhizosphere (22). In another recent study, genome-wide saturation mutagenesis of *Pseudomonas simiae* WCS417r revealed that 2% of the protein-coding genes are important for successful root colonization (23). Mutations that negatively affect rhizosphere competence, and mutations that confer a competitive advantage were identified in this study. The identification of mutations that can lead to increased root colonization (23), suggests that there is room for improvement of bacterial fitness in the rhizosphere.

In the present study we used an experimental evolution approach (24) to study how *Pseudomonas protegens* CHA0 (CHA0) evolves during repeated colonization of the rhizosphere of the model plant *Arabidopsis thaliana* (Arabidopsis). The model biological control agent CHA0 displays broad-spectrum antagonistic activity against several plant pathogenic fungi and bacteria (11), and its complete genome sequence is available (25). We performed highly controlled experimental evolution in a gnotobiotic and carbon-free sand system in which bacteria depend solely on the plant for supply of carbon. Following inoculation and establishment on the roots, bacterial populations were transferred to new plants, and this cycle was repeated eight times. We hypothesized that the repeated colonization and establishment of the bacterial population on the plant root would create an environment in which selection pressure drives the accumulation of better colonizers. *In vitro* characterization of individual bacterial colonies from these populations combined with sequencing analysis led to the identification of several evolutionary trajectories involving 35 distinct mutations that impact social traits representing inter-population communication and cooperation, carbon source utilization, motility, or biocontrol activity. By combining experimental evolution with whole genome re-sequencing, we created a powerful screen for the identification of adaptive mutations with positive effects on rhizosphere colonization.

## Results

### Mutational events in independent evolutionary lines

We previously studied five experimental evolutionary populations, referred to as lines, of CHA0 evolving in the rhizosphere of Arabidopsis in a gnotobiotic system. Independent populations were introduced on the roots and after four weeks of plant growth the populations were transferred to new plants. This cycle of transferring was repeated eight times, and we performed extensive characterizations up until cycle six to account for feasibility (26). In short, for each line, after every cycle we plated a fraction of the population on culture media and randomly picked 16 colonies for extensive phenotypic assessment of bacterial life history traits (26). In order to study adaptation at the genetic level, we selected six colonies from each line at cycles 2, 4, and 6 such that they represented most of the observed phenotypic diversity among the 16 colonies that were initially picked and characterized. These colonies, as well as six colonies from the ancestral population that was initially introduced on the roots, were re-sequenced to an average depth of 25-fold coverage (minimum 10, maximum 70) and used for the identification of single nucleotide polymorphisms (SNPs), as well as small and large insertions or deletions (INDELs). In total we thus set out to acquire genetic data for 96 bacterial colonies (5 lines * 3 cycles * 6 colonies + 6 ancestral colonies). Unfortunately, we were unable to acquire sufficient sequencing data for two colonies from line 4 at cycle 4, yielding a final set of 94 (88 evolved, 6 ancestral) re-sequenced colonies. The six ancestral colonies were all identical, indicating that there was no genetic variation in the starting population and that all observed mutations are *de novo* mutations. In total, one or more mutations were detected in 64 evolved colonies, which collectively represent 73% of the 88 characterized bacterial colonies (Table S1). We identified 5 synonymous substitutions, 20 nonsynonymous substitutions, and 4 deletions ranging in length from 1 base pair (bp) to about 400 bp, distributed over 22 genes, and 6 additional mutational events located in intergenic regions (Table 1). Mutations located in intergenic regions possibly affect transcription of nearby genes via affecting their regulatory proteins binding sites and subsequent changes in their promoter activity (27, 28). Several mutations were found to be clustered in select genes and/or regions in the CHA0 genome, *e.g.* those in the response regulator *gacA* gene (PFLCHA0_RS17965; *NC_021237:4,039,113-4,039,754*) or in the *OBC3* gene cluster involved in LPS biosynthesis (*NC_021237:2,173,707-2,196,028*)(29), but the majority of the mutations were spread across the 6.1 Mbp CHA0 genome (Figure 1). Functional characterization of the mutated genes by analyzing their Clusters of Orthologous Groups (COGs) annotation revealed that the majority of these genes are involved in transcription (COG term ‘K’), signal transduction mechanisms (COG term ‘T’), amino acid transport and metabolism (COG term ‘E’) and cell wall/membrane/envelope biogenesis (COG term ‘M’) (Figure 1, Table 1).

**Figure 1.**
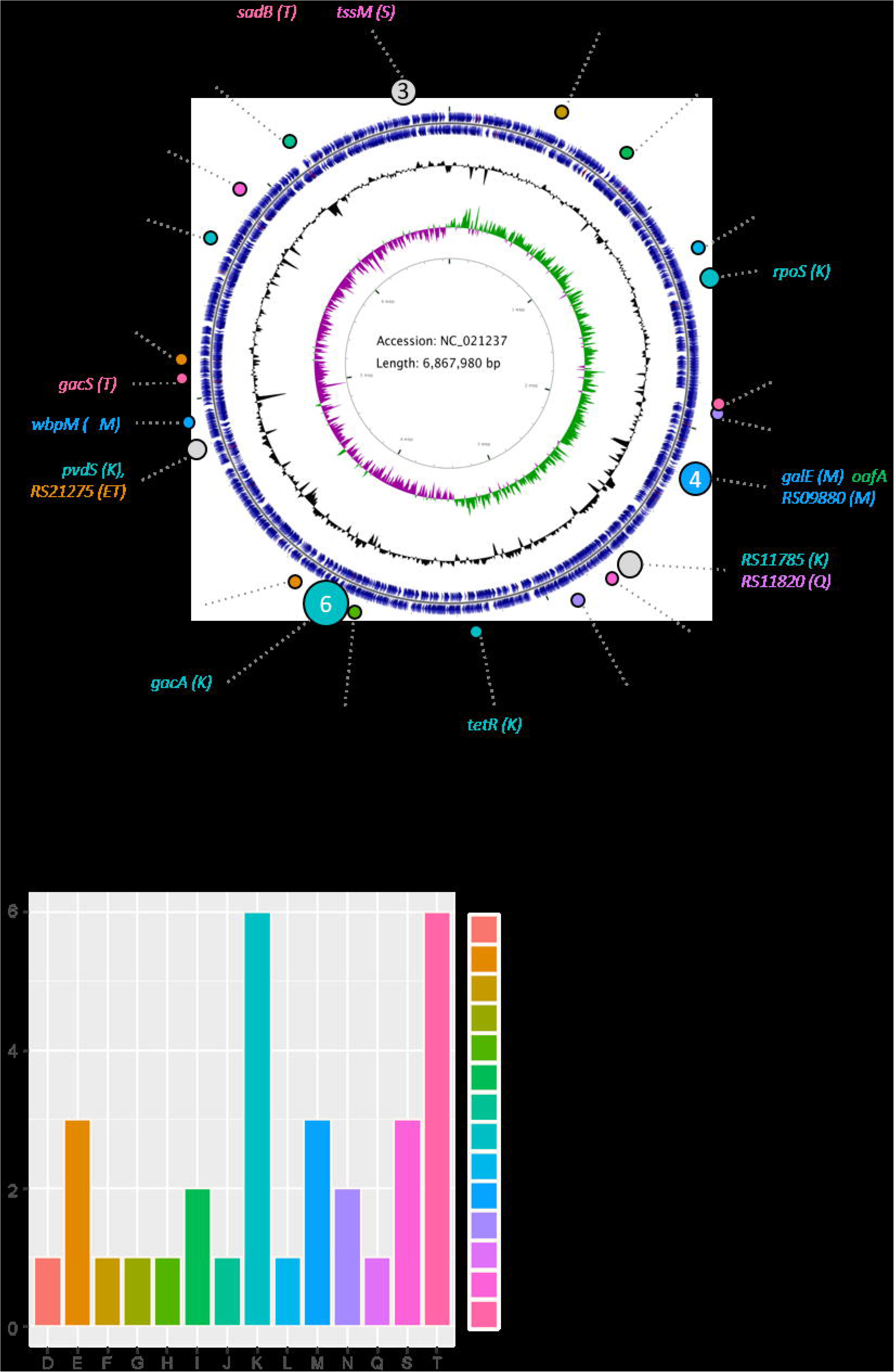
Genomic distribution of *Pseudomonas protegens* CHA0 evolutionary adaptations. **a)** Rings from inside to outside: Ring 1, nucleotide position indicator; Ring 2, green/purple GC-skew (−/+); Ring 3, %GC; Ring 4, protein-coding genes; Ring 5, distribution of identified mutations arisen during the evolutionary experiment. Functional annotations of mutated genes are indicated by color representing Cluster of orthologous groups (COGs); the key of which can be seen in panel B. **b)** Frequency of mutations per COG class, highlighting enrichment in the classes T (signal transduction), K (transcription), M (cell wall) and E (amino acid transport and metabolism).

**Table 1.**
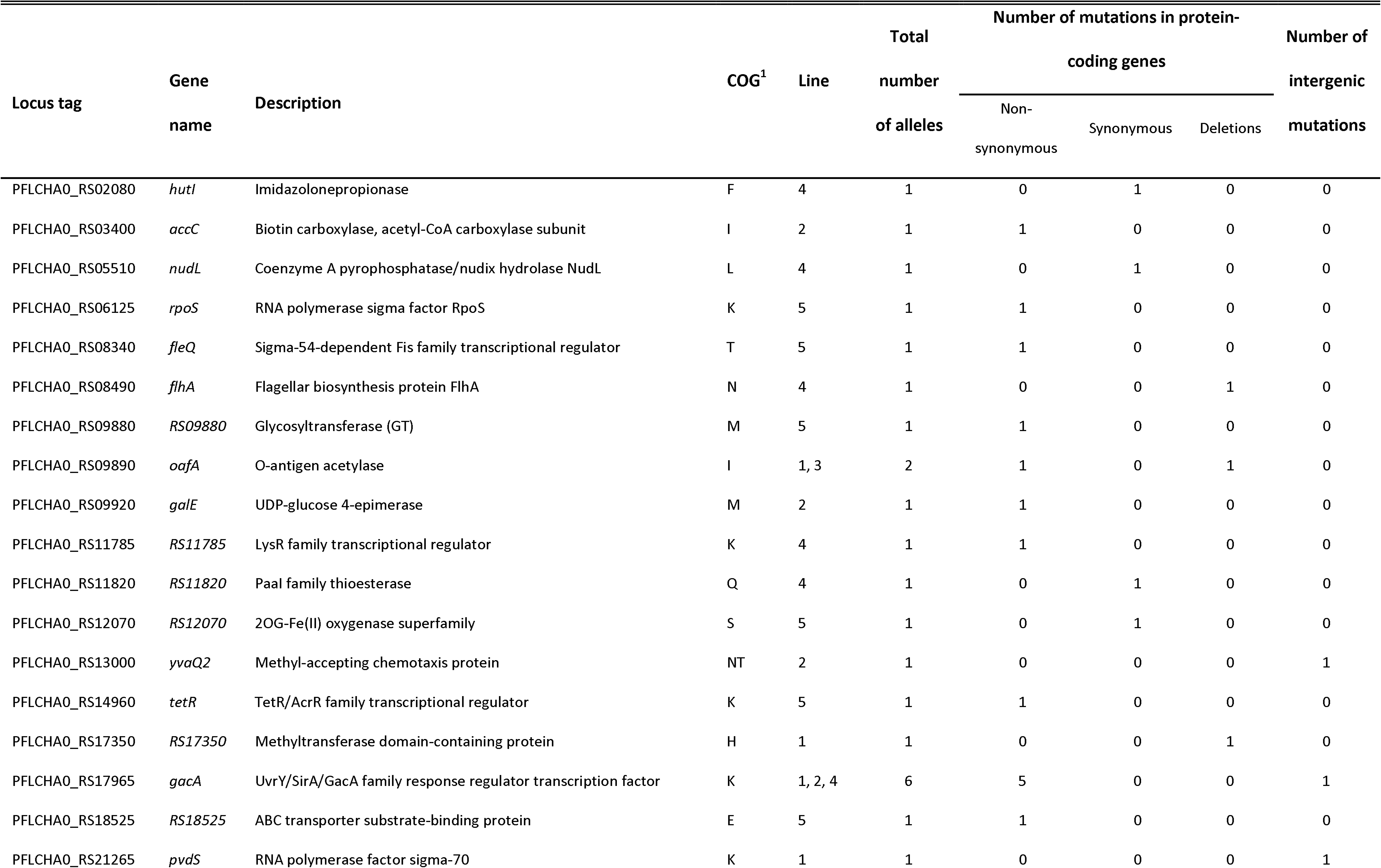
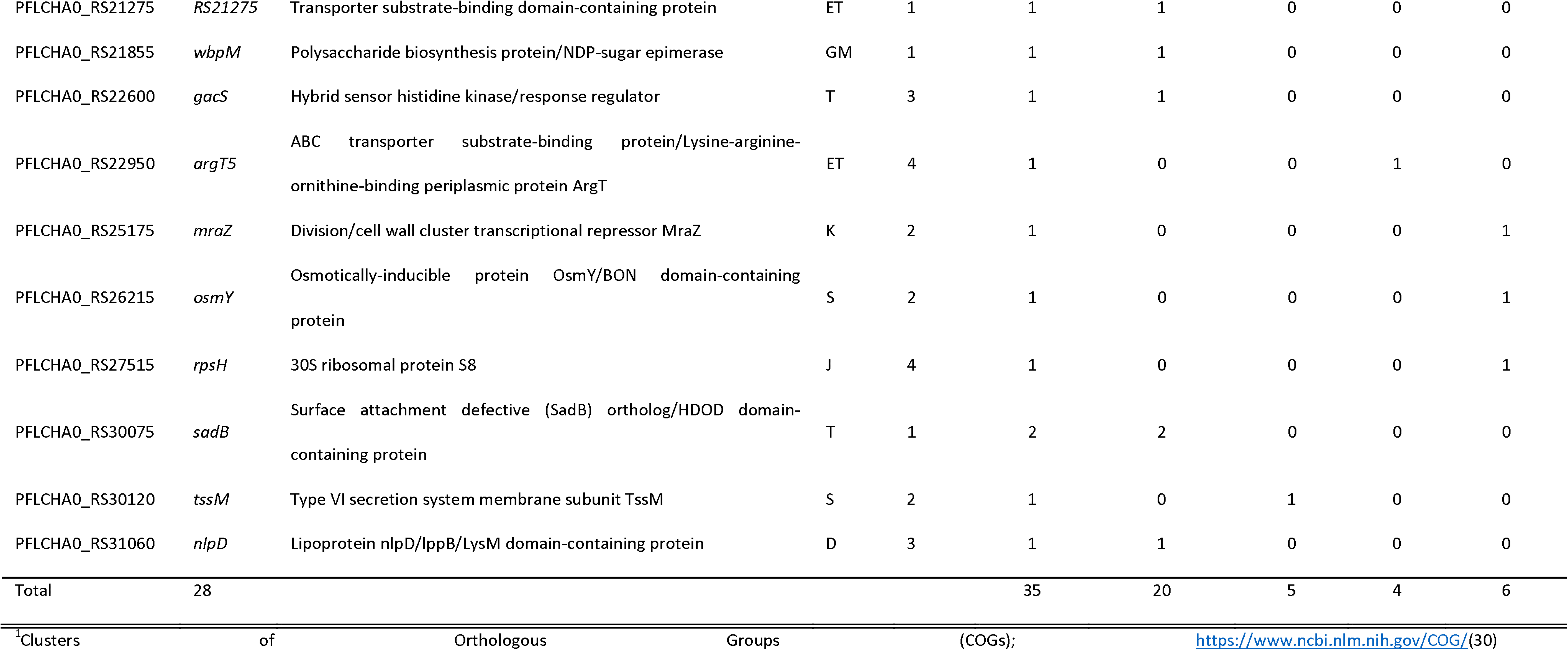
Mutations that occurred in 28 genes during Arabidopsis rhizosphere adaptation divided over five replicate CHA0 populations (lines)

### Identification of potential root colonization genes

Bacterial genes that are involved in colonization of plant roots can be revealed by identifying beneficial mutations that evolve during adaption of bacteria to the rhizosphere environment and have positive effects on root colonization. Over time, such mutants will outcompete the ancestral strain and become dominant in the bacterial population. The observation that only a limited number of mutations accumulated relative to the total number of genes within the genome across the entire experiment, makes it highly unlikely that the same gene would acquire several changes by chance in independent evolutionary lines. Nevertheless, we observe recurrent mutations in several of the same genes and/or pathways (Table 1), which is a strong indication for adaptive evolution. Genes or pathways that acquired mutations in multiple independent CHA0 populations included *gacA* and *gacS*, the *OBC3* gene cluster (*oafA*; *galE*; *PFLCHA0_RS09880* (29)), and a putative pyoverdine siderophore biosynthesis cluster (*pvdS*; *PFLCHA0_RS21275*). Other genes, like *sadB*, were targeted more than once in the same population. Because these genes were repeatedly identified in the CHA0 evolution experiment, they can be assumed to contribute significantly to bacterial fitness in the rhizosphere (Table 1, Figure 2). Moreover, it suggests that the independent evolutionary lines converge on similar evolutionary trajectories involving overlapping biological processes and molecular mechanisms.

**Figure 2.**
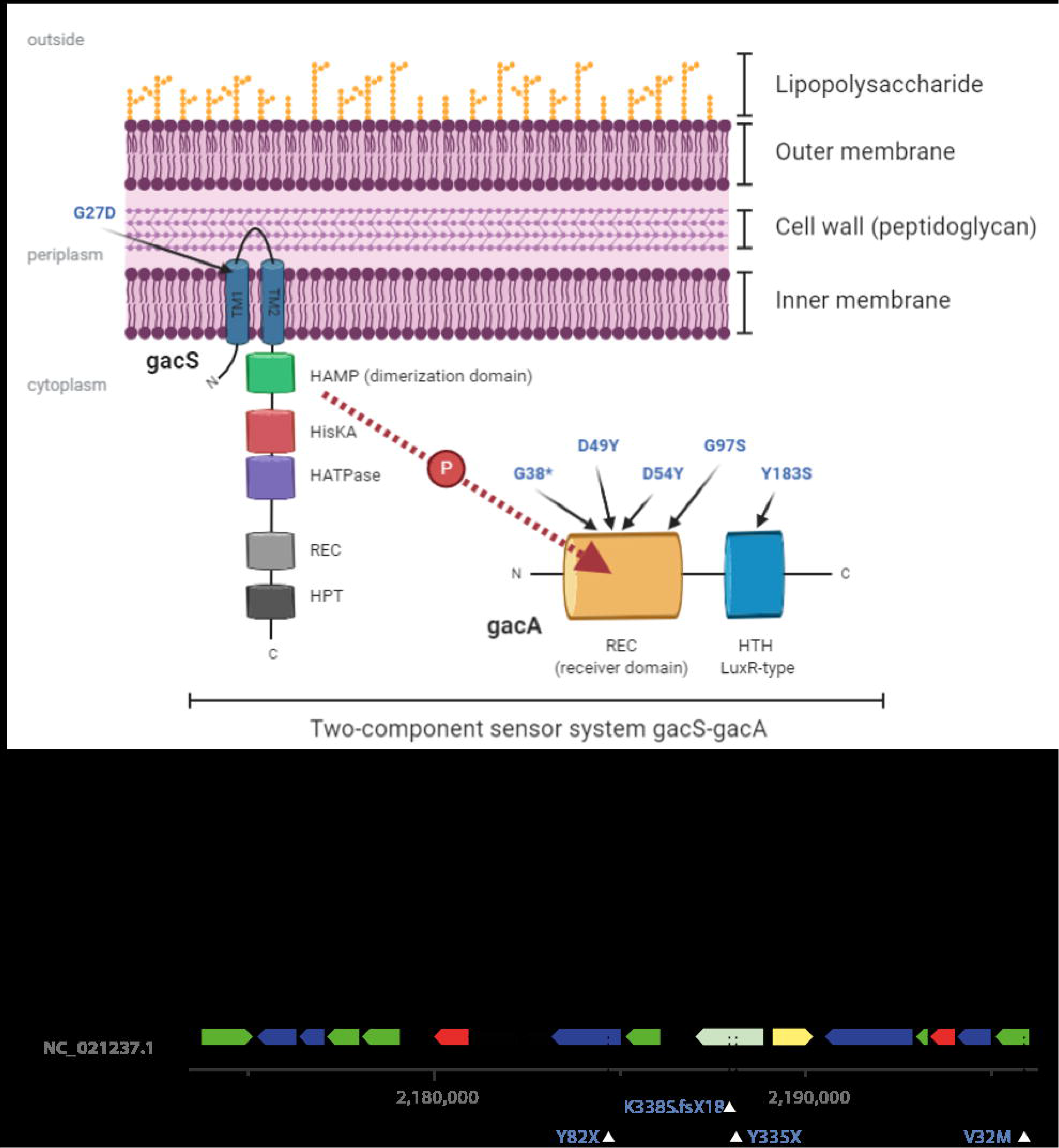
Localization of *gac* and *OBC3* mutations identified in the evolutionary experiment. **a)** Schematic view of the cell wall of the gram-negative bacterium *Pseudomonas protegens* highlighting the presence of the lipopolysaccharide (LPS) on the outside and the presence of the two-component regulator system GacS/GacA on the inner membrane and in the cytoplasm. The arrows indicate the location of the amino acid substitutions that were identified in this study in GacS (G27D) and GacA (G38X, D49Y, D54Y, G97S and Y183S). Note the travel of a phosphate group (P) from GacS to GacA upon signal perception which is accepted by either the Asp49 or Asp54 residue in GacA which are both found mutated in two different mutants. **b)** Genomic region harboring the OBC3 gene cluster responsible for the synthesis of long O-polysaccharide (O-PS) on the LPS and indicated mutations that were identified in a glycosyl transferase (RS09880; Y82X), the O-antigen acetyltransferase oafA (K338S.fsX18 and Y335X) and the UDP-glucose 4-epimerase (galE; V32M).

To illustrate the diverse evolutionary trajectories, we constructed phylogenetic trees of each replicate population (line) using all mutations that were detected (Figure 3). In each line, both the number and depth of branches are very different (Figure 3, Table 2). In line 1 and 3, the populations were swept early by *oafA* mutants and later by specific *gac* mutants, while in line 2 and 4, co-existence of multiple genotypes characterized by *gac* mutations was observed indicative of clonal interference between lineages (Figure 3; (31)). Notably, the relative abundance of each genotype in the population is estimated from a small set of sequenced individuals and therefore is expected to lack precision. The evolved isolates contain one to four mutations in general, and in each line, 3 (line 3) to 9 (line 2 and 4) mutations were identified in total. All mutations were unique for each line, as is expected for independently evolving populations.

**Figure 3.**
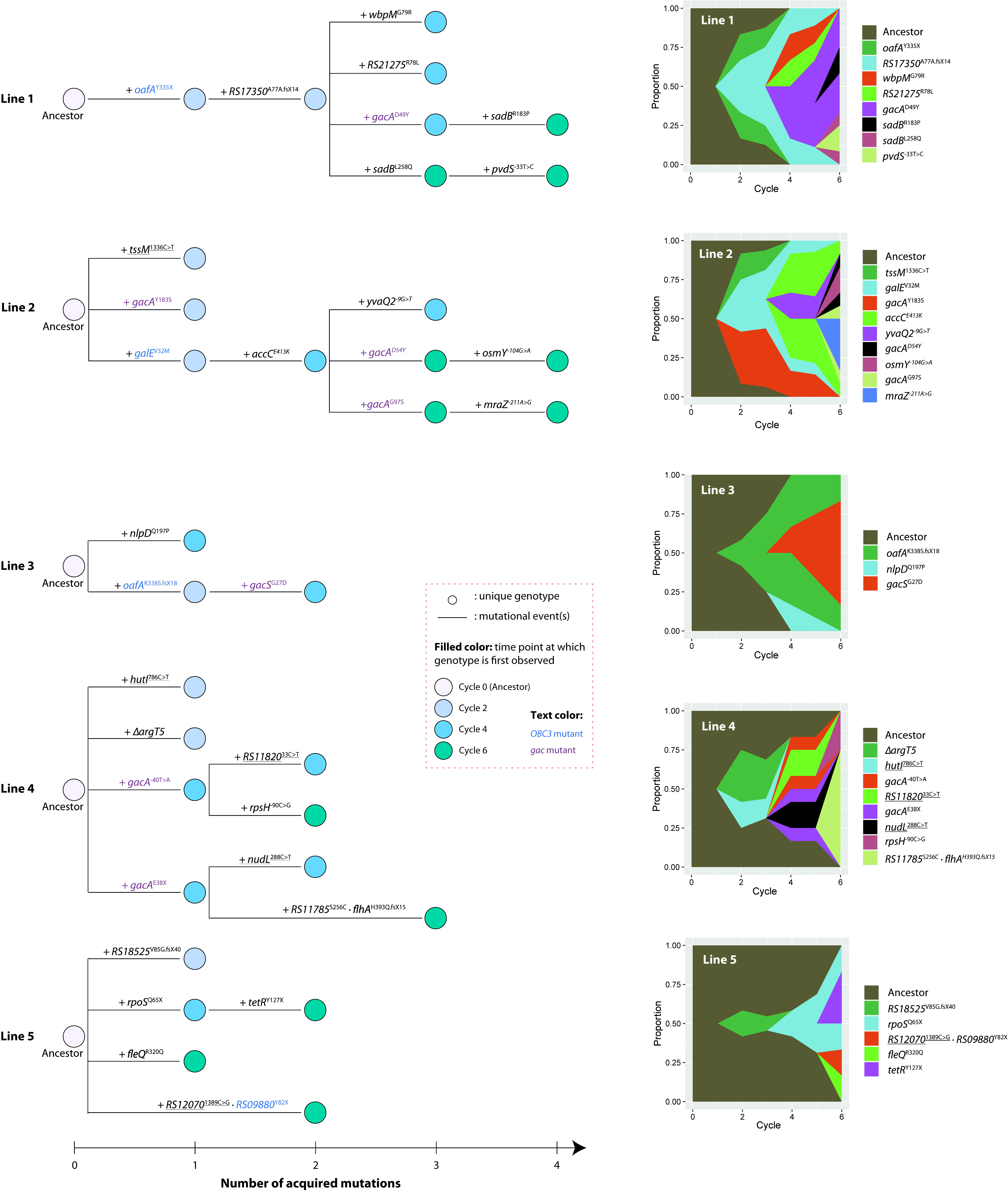
Phylogenetic trees and Muller plots of five independently evolving *P. protegens* CHA0 populations. Left) Phylogenies for 18 genomes from each population (16 for the population in experimental line 4), based on the sequential appearance of mutations are shown. Synonymous mutations are underlined. Circles represent unique bacterial genotypes. The color of the circle fill represents the time point at which a genotype was firstly detected. *OBC3* and *gac* mutants are highlighted in blue and purple, respectively. Right) Muller plots depicting the dynamics of mutant alleles during the evolutionary experiment. The Muller plots show the estimated frequencies, by their height, of 33 mutations in the respective population over 6 experimental cycles. Descendant genotypes are shown emerging from inside their respective ancestors. The frequency of each mutation in the respective population can also be found in Table 2. Muller plots are prepared using *ggmuller* in R (101).

**Table 2.**
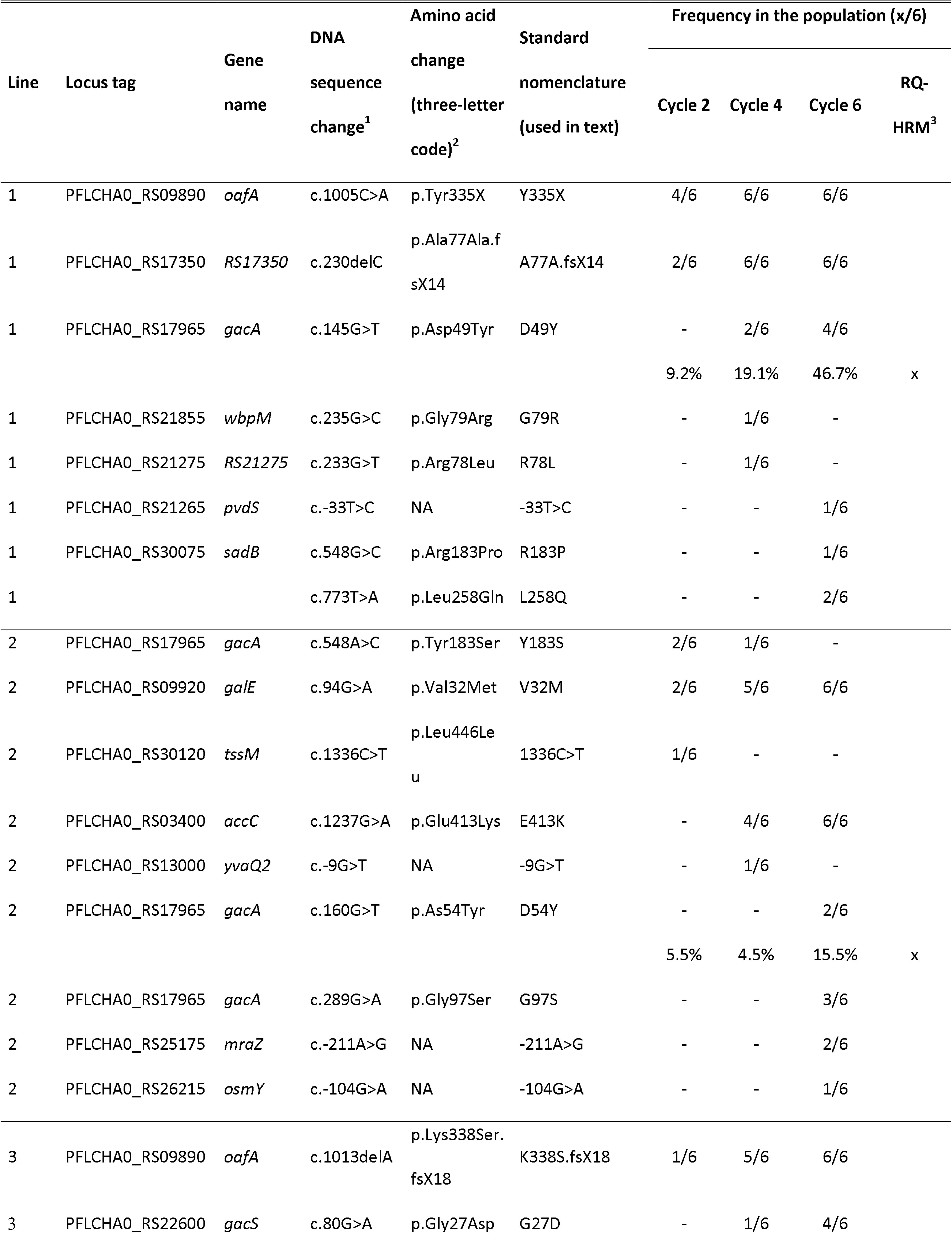
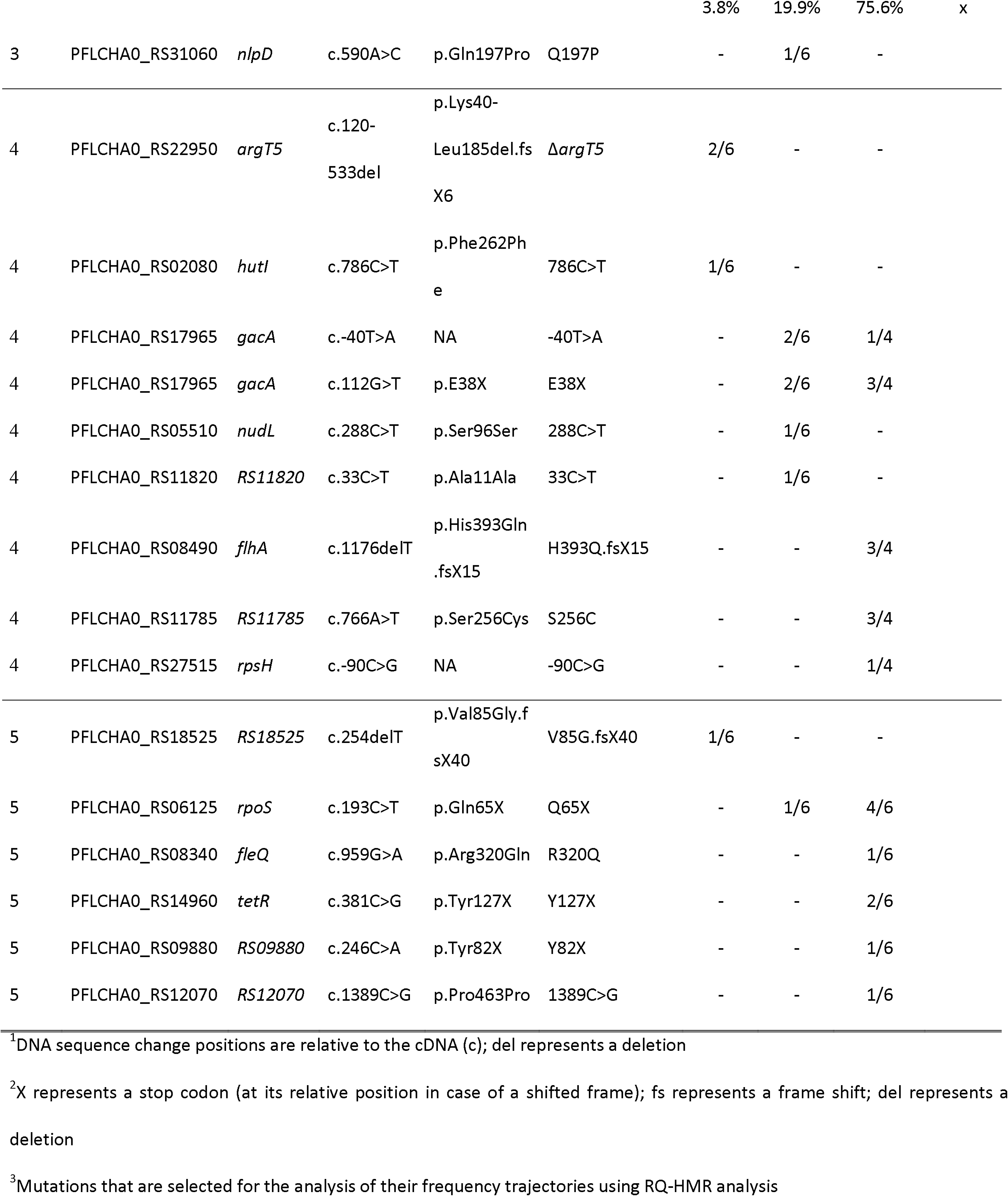
Nomenclature for CHA0 mutations and variants and their frequency/cycle in each respective population.

As the frequency of each mutation (Figure 3, Table 2) was determined from only six bacterial colonies that were isolated and sequenced from each evolutionary line at cycles 2, 4 and 6, we further investigated and accurately measured the population-level frequency at the end of each experimental cycle. We determined the frequency of three *gac* mutations, *i.e.*, *gacA* ^D49Y^*, gacA*^D54Y^, and *gacS*^G27D^, with increased accuracy, *i.e.* by PCR-based high-resolution melting (HRM) analysis incorporating mutation-specific HRM probes (Table S2), and increased sampling depth, *i.e.*, at the end of experimental cycles 1 to 8. The HRM methodology allows for the accurate quantification of mutant frequency across a wide range for all three mutations (*p*<0.001; Figure S1). Using this method, we found that the population-level frequency of these three mutations in the respective evolutionary lines corresponded remarkably well with the frequency previously obtained from the cultured and sequenced isolates (Table 2, Figure S2). These findings corroborate the culture-based quantification of mutant frequency suggesting they provide a reasonable measure for population-level frequency. Intriguingly, it can be seen that the *gacS*^G27D^ mutation reaches fixation after eight experimental cycles, while the *gacA*^D49Y^ and *gacA*^D54Y^ mutants stabilize at around 50% and 25%, respectively (Figure S2). Stabilization of mutations is indicative of frequency-dependent (FD) selection putatively reinforced by clonal interference with co-existing lineages carrying beneficial mutations in *sadB* (L258Q) and *gacA* (G97S) (Figure 3). FD selection describes the phenomenon that the fitness of a particular genotype or phenotype is dependent on its frequency. Such context-dependency has been linked to cheating behaviour in which microbial cells that lack or have limited production of certain costly compounds benefit from other cells that do produce these compounds. When a minimal amount of such compound increases genotype or phenotype fitness, FD selection can occur resulting in stabilization of the mutation frequency.

### Early adaptations are driven by cell surface-related genes

In general, mutations that are fixed early on in the rhizosphere adaptation process tend to have a high selective advantage (32). Disruptive mutations in *oafA*, resulting in premature stops halfway the coding region, were detected as the first acquired mutations in two independent evolutionary lines (lines 1 and 3) and appear to have swept the population in the following generations (Table 2). *OafA* is part of the O-polysaccharide (O-PS, O-antigen) biosynthesis cluster 3 (*OBC3*)(29), and encodes an O-acetyltransferase which is postulated to acetylate the O-antigen component of the outer membrane LPS (33). Another *OBC3* mutation that accumulated early on in the rhizosphere adaptation process, *galE*^c.94G>A^, leads to an amino acid substitution (V32M) in galE and this mutation swept through the population in evolutionary line 2, reaching fixation in cycle 6 (Table 2; Figure 3)*. GalE* encodes an UDP-glucose 4-epimerase which is involved in O-antigen and LPS core biosynthesis (34–36). One colony with a mutation in a third *OBC3* cluster gene, *RS09880*, encoding a putative glycosyl transferase (GT) was found in cycle 6 of evolutionary line 5. Thus, in four out of the five evolutionary lines mutations that likely affect bacterial LPS structure appeared during rhizosphere adaptation, and these mutations became dominant in three out of four evolutionary lines. These results strongly suggest that modifying bacterial cell surface structure is an important bacterial strategy in early adaptation to the rhizosphere.

### Adaption driven by global regulators

In the present study, six mutations were detected in *gacA* in three out of five evolutionary lines, representing approximately 20% of all missense mutations. Notably, in evolutionary lines 2 and 4, multiple *gacA* alleles accumulated, some of which were detected in early experimental cycles (Table 2). Additionally, a *gacS* mutation accumulated in line 3. *GacA* and *gacS* encode the main constituents of the conserved GacA/GacS two-component regulator system, *i.e.*, the hybrid sensor histidine kinase GacS and the cognate response regulator GacA (Figure 2). In gram-negative bacteria, activation of GacS results in cross-phosphorylation of GacA via phosphotransfer which in turn leads to activation of the expression of the small RNA genes *rsmY* and *rsmZ* via its helix-turn-helix (HTH) domain-binding domain (37). In CHA0, this regulatory pathway is known to control quorum sensing as well as secondary metabolism and stress resistance (38–41). In the closely related strain *P. protegens* Pf-5 this pathway was shown to have a big impact on bacterial genome-wide gene expression, affecting the expression of more than 10% of the annotated genes (42). Similarly, *gacA* mutants that arose on the edge of swarming Pf-5 colonies showed dramatically altered genome-wide gene expression patterns (43).

Including *gacS* and *gacA*, about half of the mutated genes in this study are global regulators or sigma factors (Table 1, Figure 1). This high frequency suggests that global regulator-controlled networks are evolvable and play a major role in rapid bacterial adaptation. Pleiotropic adaptive mutations in global regulator genes have been shown to be important for bacterial adaption both in laboratory (44, 45), and in natural settings (46, 47). Remodeling and continuous optimization of existing regulatory networks by single mutations is an important strategy for bacterial adaptation to the host (48).

### Bacterial motility

Bacterial motility is an important trait for rhizosphere competence, mediating colonization of distal parts of the root system (49), and both LPS and the GacS/GacA two-component regulator system are known to affect this trait. The O-antigen side chain of the LPS was reported to contribute to swimming and swarming motility in the plant pathogenic bacterium *Erwinia amylovora* (50, 51). The GacS/GacA two-component regulator system controls bacterial motility, for example by affecting transcription of genes related to flagella and biosurfactant biosynthesis (43, 49, 52). We also identified several other mutations across the various evolutionary lines that can be linked to bacterial motility. For instance, a disruptive mutation in the flagellar biosynthesis protein-coding gene *flhA* (H393Q.fsX15) that is involved in the biogenesis of the bacterial flagellum (Table 1) appeared in evolutionary line 4, reaching up to a predicted frequency of 75% (Table 2; Figure 3). In *P. fluorescens* Pf-5, and in *Pseudomonas aeruginosa* PA01 FlhA is reported to be essential for swimming motility (42).

Furthermore, we identified one amino-acid substitution, R320Q, in FleQ and two in SadB, R183P and L258Q respectively, that based on sequence similarity to well-studied, homologous proteins in other bacteria can be linked to motility in addition to several other bacterial traits. FleQ is a σ^54^-dependent Fis family transcriptional regulator which regulates flagellar motility, biofilm formation as well as Pel exopolysaccharide (EPS) production in response to cellular c-di-GMP levels in *P. aeruginosa* (53). *P. protegens* FleQ shares 84% sequence identity and 98% sequence coverage with *P. aeruginosa* FleQ and like *P. aeruginosa* FleQ it is comprised of the N-terminal flagellar regulatory FleQ domain (PF06490), a central AAA+/ATPase σ^54^ –interaction domain (PF00158), and a C-terminal Fis-type HTH DNA-binding domain (PF02954). The R320Q substitution we identified here is found in the AAA+/ATPase σ^54^-interacting domain in between the arginine (Arg) finger (300-303) and a c-di-GMP-binding motif, ExxxR (330-334) (53). The conserved arginine residues in FleQ, including the here mutated Arg^320^, are thought to be important for protein oligomerization, and substitution of any of these residues abolishes ATPase activity in *Vibrio cholerae* EpsE completely (54). Finally, *sadB*, encodes a HD-related output domain (HDOD)-containing protein (Table 1) that shares 77% sequence identity and 99% sequence coverage with *P. aeruginosa* SadB. In *P. aeruginosa*, SadB stimulates Pel EPS production and the chemotaxis-like cluster CheIV, which in turn affect flagellar motility as well as biofilm formation (55). In *Pseudomonas fluorescens* F113, SadB, together with FleQ control flagellar motility, dependent and independent of the GacS/GacA two-component regulator system (56, 57).

Parallelism of targeted mutations on this functional motility pathway impelled us to assess bacterial motility and track its dynamics across all evolutionary lines. We selected all *OBC3* and *gac* mutants as well as mutants in *sadB*, *fleQ* and *flhA*. Additionally, we included two *gacA* mutant progenitors with mutations in *accC* and *RS17350*, encoding a biotin carboxylase and a methyltransferase, respectively, plus two *gacA* mutant descendants with mutations in *osmY* and *mraZ* that encode for an osmotically-inducible protein and a transcriptional repressor, respectively (Table 1). Altogether, we assessed swimming and swarming motility of seventeen distinct genotypes (Figure 4; Figure S3). We found that *OBC3* mutants, the *gacA* progenitors with mutations in *accC* and *RS17350*, and the *gacA* descendants with mutations in *osmY* and *mraZ* were unaltered compared to their respective ancestors when considering both swimming and swarming motility, with the exception of a small yet significant increase in swarming motility in *oafA*^Y335X^. However, g*ac* mutants themselves were significantly affected; both in swimming, which is generally enhanced, and in swarming, which is repeatedly decreased (Figure 4). *SadB* mutants, like *gac* mutants display enhanced swimming and worsened swarming when compared to their respective progenitors. Oddly, both the *fleQ* and the *flhA* mutant displayed severely reduced swimming and swarming motility, thus representing two examples of an alternative evolutionary route towards adaptation in the rhizosphere in which motility is reduced. Loss of motility might coincide with another trait in these cases, such as EPS production and/or biofilm formation, regulated via shared yet oftentimes opposing mechanisms. Also, the frequency of the *fleQ* mutant is low, *i.e.*, only one out of six isolates from cycle 6 of experimental line 5 carried this mutation and therefore whether this mutant is truly beneficial is unclear. On the other hand, the *flhA* mutant was found in three out of the four cultured and sequenced isolates from cycle 6 of line 4 and therefore could represent a significant proportion of the total population.

**Figure 4.**
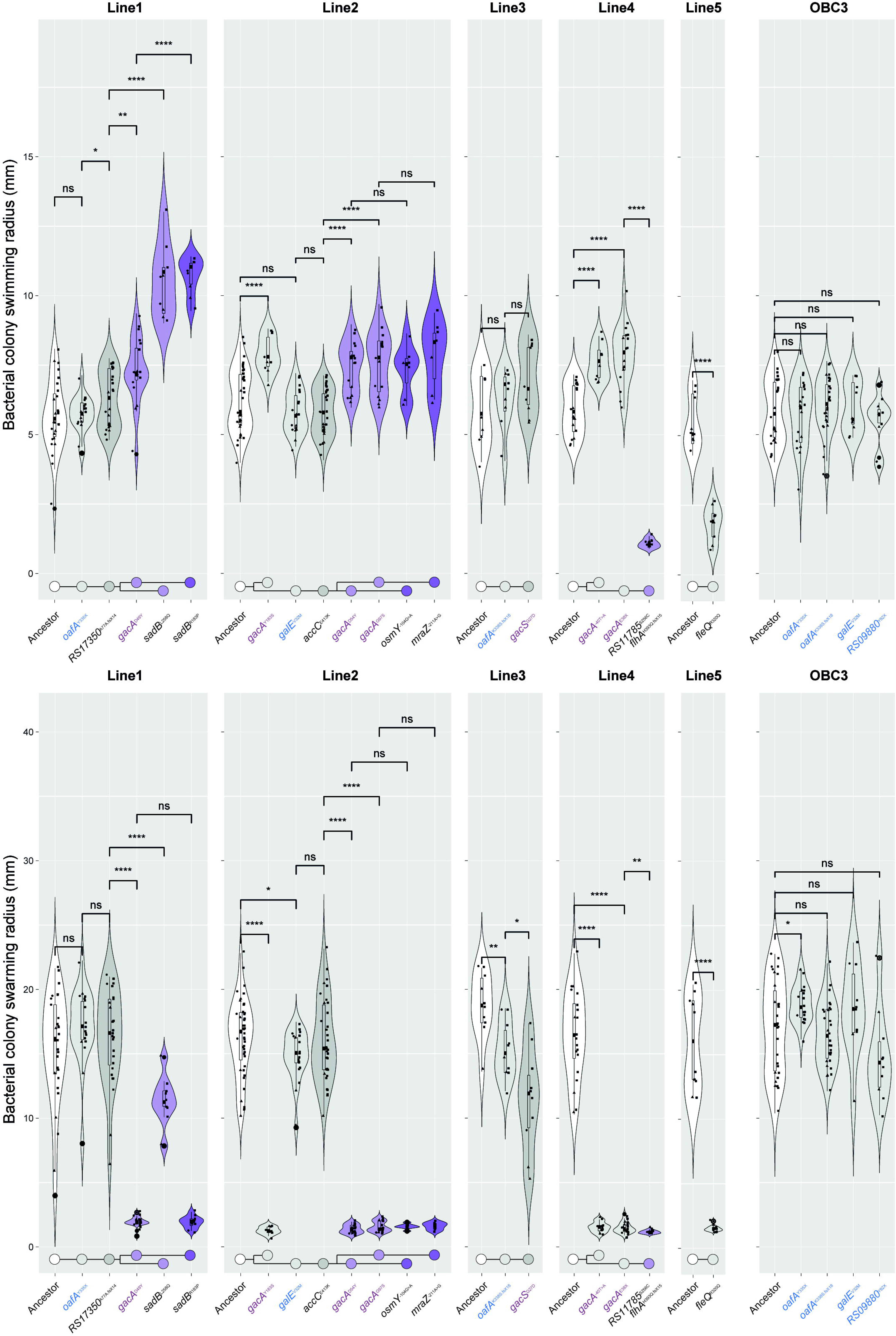
Evolved, root-competent mutant *P. protegens* CHA0 strains are characterized by enhanced swimming and impaired swarming. The accumulative effects of parallel and sequential mutations on swimming motility (top panel) and swarming motility (bottom panel) are shown. For each experimental line, strains carrying mutations that can be connected to bacterial motility, or that are the ancestor or descendants of such mutant strains, were studied using typical swimming and swarming assays on Cook’s Cytophaga (CC) medium (95). The colony area as a measure for bacterial motility was determined using ImageJ. The combined data from three independent experiments is shown, and datapoints from each experiment can be discerned by their respective shape. The sample size, *n*, varies per genotype as related strains are combined on each plate, and we report the minimum and maximum *n* per replicate experiment. For swimming assays, circles represent replicate 1 (2≤n≤6) (excludes *osmY*^−104G>A^ and *mraZ*^211A>G^), triangles replicate 2 (3≤*n*≤15), and squares replicate 3 (4≤n≤20). For swarming assays, circles represent replicate 1 (1≤*n*≤3) (excludes *osmY*^−104G>A^ and *mraZ*^−211A>G^), triangles replicate 2 (2≤*n*≤6), and squares replicate 3 (7≤*n*≤35). Obvious outliers were removed after pooling. Significant differences in motility between a genotype and its respective progenitor were determined by unpaired t-test analysis (swimming, 7≤*n*≤41; swarming, 7≤*n*≤44; *α=0.05, **α=0.01, ***α=0.005, ****α=0.001, ns = non-significant) and the result is shown above each comparison. The genealogy of the mutations is shown below each experimental line highlighting both parallel (branching) and sequential mutations. Colours depict the number of acquired mutations relative to the ancestor (white), and these are light-grey: 1, grey: 2, light-purple:3 and purple: 4.

### Dynamics of global phenotypic change

Since natural selection eventually operates at the phenotypic level, revealing bacterial global phenotypic evolutionary dynamics can help us to identify traits that are under selection. Moreover, beneficial genetic mutations can be predicted if they are linked to well-known root colonization traits. A broad range of 30 bacterial traits including different aspects of bacterial life-history traits, were examined for the sequenced isolates, which allows for genotype-phenotype association analysis.

As shown in Figure 5a, the 30 bacterial traits separated into four clusters that share a similar pattern across the different mutant genotypes and this clustering is supported by model-based clustering analysis (Figure S4). Growth of the bacteria in 1/3 strength King’s B (KB) medium was positively correlated with siderophore and exoprotease production, tryptophan side oxidase activity, and growth inhibition of the bacterial plant pathogen *Ralstonia solanacearum*. Thus, this cluster, designated cluster 1, contains traits associated with bacterial social behavior, related to microbe-microbe communication and cooperation, such as the production of public goods. In a principal component analysis (Figure S5a), the first principal component (PC1) is strongly correlated with all five traits, with a total explanation of 64.5% for all variables. Cluster 2 contains traits linked to carbon source utilization. For this cluster PC1 is strongly correlated with all carbon source usage-related traits, with a total explanation of 83.9% for all variables (Figure S5b). A third cluster was observed for the bacterial ability to form a biofilm, to produce indole-3-acetic acid (IAA), and to inhibit the growth of two fungal plant pathogens, with a total explanation of 82.4% by PC1 (Figure S5c). Finally, cluster 4 contains all seven traits that are related to bacterial resistance to biotic and abiotic stresses. The first principal component (PC1) is strongly correlated with all seven traits, with a total explanation of 53.9% for all variables (Figure S5d). For these four clusters, the PC1 (or -PC1) value is used as a proxy to present the general performance of all the traits that clustered together.

**Figure 5.**
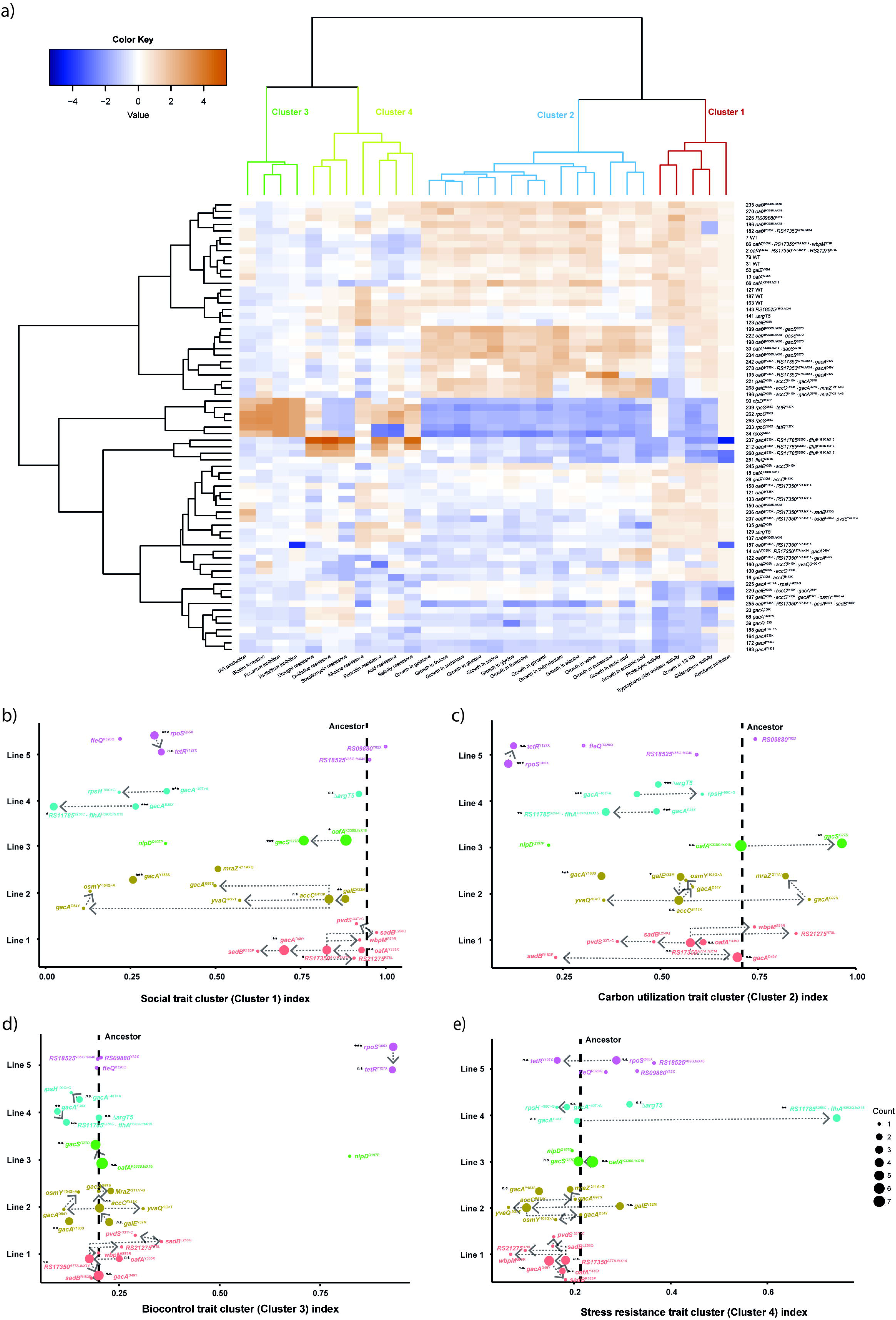
Genotype-phenotype association analysis of evolved bacterial populations. Isolates without mutations or only with synonymous mutations were excluded for this association analysis **a)** Heatmap representing the performance of 30 life-history traits of six ancestral and 63 evolved isolates. The “ward.D2” method was used for hierarchical clustering, based on the dissimilarities of the set of objects being clustered. The four clusters depicted on top are model-predicted. Each column illustrates one bacterial life-history trait, indicated on the bottom of the figure. Each row represents an isolate, indicated by its sample ID number and mutational events. **b-e)** Principal component analysis was applied to generate a general symbolic index for each model predicted cluster. The proxy values of each cluster were normalized, higher values represent better performance. To illustrate better the accumulative effects of each mutational event, ANOVA was applied to reveal the phenotypic change between each genotype and their identified nearest ancestor in each line (only applied for genotypes that have been detected more than once; n>=2). Asterisks alongside the mutational events indicate significant differences (*α=0.05, **α=0.01, ***α=0.001; n.s. = non-significant). Colour-filled circles represent bacterial genotypes, with different colours representing independent evolving populations and the size denotes the number of replicates (n). The dashed line represents the average value of the ancestor.

As is shown in Figure 5b, all five evolutionary lines showed a parallel trend of accumulative decline of social traits. Mutations in *gacA, gacS,* and *rpoS* resulted in significantly decreased bacterial social traits, and this was also observed for the double mutant *RS11785*^S256C^ *· flhA*^H393Q.fsX15^ relative to the background *gacA*^E38X^ mutation. *RS11785*, like *gacA*, encodes a LysR-type transcriptional regulator. In addition to these global regulators, earlier mutations in *oafA*, *galE* and *RS17350*^A77A.fsX14^encoding a putative methyltransferase, also resulted in a significant but relatively small decrease in social traits. This parallel decline of the bacterial social traits index suggests a negative selection of bacterial social behavior, especially the production of costly public goods such as siderophores and exoproteases.

All traits related to the utilization of 14 different carbon sources, that were selected based on their reported presence in root exudates of Arabidopsis (58) grouped in one cluster (cluster 2; Figure 5a). This suggests that carbon source utilization is co-regulated. Different mutations in *gacS* and *gacA* resulted in contrasting bacterial carbon source utilization. A mutation in the first N-terminal transmembrane domain of GacS (G27D) (Figure 2) resulted in significant enhancement of carbon source utilization (Figure 5a, c). A similar trend was observed for mutant *gacA*^G97S^. In contrast, the majority of evolved genotypes, including most mutations in *gacA* and *rpoS*, showed a reduced ability to utilize carbon sources (Figure 5a, c).

In contrast to the previous two clusters, traits in clusters 3 (Figure 5d) and 4 (Figure 5e) were more stable, as most genotypes behave like the ancestor. Only two *gacA* mutants, *gacA*^E38X^ and *gacA*^Y183S^, were associated with a significant decline of bacterial traits associated with biocontrol, *i.e.*, antifungal activity and biofilm formation, while the *rpoS*^Q65X^ mutation resulted in a significant increase for these traits. The double mutation *RS11785*^S256C^ · *flhA*^H393Q.fsX15^, in the background of the disruptive *gacA*^E38X^ mutation (Figure 5e), was the only mutational event that led to a significant increase of general resistance to various environmental stresses. *FlhA* encodes the flagellar biosynthesis protein FlhA and is linked to bacterial motility. The LysR-type transcriptional regulator RS11785 comprises a N-terminal HTH DNA-binding domain (PF00126) and a C-terminal substrate binding domain (PF03466). This substrate binding domain is linked to aromatic compound degradation and resembles that of the type 2 periplasmic binding proteins or PBP2s. PBP2s are responsible for binding and uptake of several substrates including polysaccharides and several amino acids. We used HHpred (59) for protein remote homology detection and three-dimensional structure analysis to assess the possible consequence of the S256C amino acid substitution. This analysis identified OxyR from *Escherichia coli* as the best template for modeling (E-value: 2.9e-30, Score: 166.45). OxyR represents the master peroxide sensor in gram-negative bacteria, which operates via intramolecular disulfide-bond formation in the regulatory PBP2-like domain. RS11785 is unlikely to represent the master peroxide sensor in *P. protegens* CHA0 because another CHA0 protein encoded by *PFLCHA0_RS030065* is much more similar to OxyR (89% percent pairwise sequence identity versus 29% for RS11785). Nevertheless, it is tempting to speculate that the serine (S, Ser) to cysteine (C, Cys) amino acid substitution (S256C) we observed here might influence regulatory activity by altered intramolecular disulfide-bond formation as such bonds are governed by pairs of cysteine residues. Lastly, RS11785 bears resemblance (37% sequence identity, 93% sequence coverage) to the LysR-type PBP2-domain containing regulator alsR which regulates the activity of the acetoin operon (*alsSD*) in response to several signals such as glucose and acetate in *Bacillus amyloliquefaciens*. Acetoin (3-hydroxy-2-butanone, a precursor for 2,3-butanediol) can elicit induced systemic resistance (60), and is linked to general plant growth promotion in *Bacilli*. In the absence of singular mutations in *flhA* and *RS11785*, the molecular mechanism underlying the observed enhanced environmental stress resistance in this double mutant remains to be clarified but we hypothesize that altered regulatory activity by RS11785^256C^ is the most likely cause.

## Discussion

The rhizosphere is a nutrient-rich environment for root-associated bacteria. However, to access the available nutrients bacteria must overcome various challenges, including the plant immune system, presence of competing and/or predatory microorganisms, and abiotic stresses. In the gnotobiotic binary system used in this study we tracked changes in *P. protegens* CHA0 in the rhizosphere of Arabidopsis under reproducible and controlled conditions without interference of complex interactions with other microbes. Mutations affecting global regulators, bacterial cell surface structure, and motility accumulated in parallel in our evolutionary experiment, revealing at least three important strategies of bacterial adaptation to the rhizosphere.

### Global regulators and rhizosphere adaptation

The GacS/GacA two-component regulator system controls the production of antimicrobial secondary metabolites, exoenzymes, and siderophores, but also biofilm formation, stress responses, motility, and quorum sensing (13, 39, 42, 61, 62). In the present study, mutations in the GacS/GacA two-component regulator system caused dramatic changes in several bacterial phenotypic traits, including motility, carbon source utilization, and social traits (Figures 4 and 5). The overall gain in swimming motility is not unexpected, as swimming motility, driven by the flagellum apparatus, has been repeatedly reported to be an important root colonization trait in root-associated bacteria, including several *Pseudomonas* spp. (17, 63–65). In line with this, genome-wide transposon disruption mutant analysis by Cole and coworkers (2018) showed that the majority of motility-related genes in *Pseudomonas simiae* WCS417 are positivelyassociated with Arabidopsis root colonization (23).

In contrast with the observed enhanced swimming motility across the evolutionary selection lines, swarming motility was found to be severely hampered throughout, and this appears to be the case especially for *gac* mutants. Swarming, like swimming, is driven by the flagellum but in addition depends on the production of several compounds, including quorum sensing molecules and biosurfactants. *P. protegens* CHA0 is a known producer of the biosurfactant orfamide A and the GacS/GacA two-component system is known to be an important regulator for its biosynthesis (66). These results fit observations by Song and co-workers (43) in the closely related *P. protegens* strain Pf-5. During swarming motility, *gac* mutants emerged that lack production of the surfactant orfamide. These mutants cannot swarm, but they co-swarm with orfamide producing cells (43).

Remarkably, we also identified two completely non-motile mutants in our experiments with disruptions in the *fleQ* and *flhA* genes. Possibly, these mutants adhere much better to the root surface or might be able to form a biofilm more rapidly or strongly as these traits are often inversely correlated with bacterial motility (67–69). Future experiments should reveal whether such trade-off between bacterial motility and root adherence indeed underlies our observations in this evolutionary experiment.

The decreased production of public goods, such as siderophores and exoproteases, which were observed in all independent evolutionary lines (Figure 5), could be beneficial for bacterial fitness by saving energy and primary metabolites. For example, adaptation of *Pseudomonas aeruginosa* to the human host through mutations in regulators was accompanied by loss of siderophore production, secreted proteases, and biofilm formation (70, 71).

### Cell surface structures as bacterial adaptation targets

Bacterial cell surface components are the first line of defense against environmental stress and interplay with hosts (72–74). LPS is a central outer membrane component for gram-negative bacteria, and exhibits structural adaptability that is contributed especially by its O-antigen part (72, 74). Bacterial O-antigen structure modification plays an important role in evasion of host immunity (75), and has the potential to change host-bacteria interactions (72, 74, 76). In plant pathogenic bacteria, LPS components are important virulence determinants (77, 78), which can activate a variety of defense-related responses (77, 79–81). In *P. fluorescens* the O-antigen component has been implicated to induce systemic resistance in radish (82) and Arabidopsis (83).

We observed parallel mutations in genes that are involved in LPS biosynthesis and structure modification. In three out of the five evolutionary lines, the first fixed mutations were identified in *oafA* and *galE* that are annotated as O-antigen biosynthesis and structure modification related (Table 1). *OafA* encodes an O-acetyltransferase, which is postulated to modify the O-antigen by acetylation (33). The enzyme GalE, UDP-galactose 4-epimerase, is involved in the interconversion of UDP-glucose to UDP-galactose, an essential intermediate for LPS core and O-antigen structures (34–36). Inactivation of *galE* in *Porphyromonas gingivalis* resulted in shortening of the O-antigen (84), and in *Bradyrhizobium japonicum*, disruption of *galE* resulted in the complete absence of O-antigen (34). Thus, it is tempting to speculate that evasion of the plant’s immune response plays a role in adaptation of CHA0 to the rhizosphere of Arabidopsis.

In conclusion, the observed bacterial genetic and phenotypic adaption dynamics emphasize important roles for global regulators, motility, and cell surface structure in bacterial adaptation to its host. The parallel emergence of mutations in similar genes resulted in specific fitness advantages for mutants in the rhizosphere, suggesting that this evolutionary process is driven by the rhizosphere environment.

## Materials and Methods

### Experimental setup

We set up an experimental evolution experiment with *Arabidopsis thaliana* (Arabidopsis) ecotype Col-0 as host plant and *Pseudomonas protegens* CHA0 (CHA0) as the evolving bacterial strain. CHA0 (85) is a model strain originally isolated from roots of tobacco plants grown in soil naturally suppressive to black root rot (86). CHA0 was chromosomally tagged with GFP and a kanamycin resistance cassette (Jousset et al 2006) to enable consistent tracking of the strain and identification of contaminations. We previously described the setup of the evolutionary experiment in great detail (26). In brief: the ancestral bacterial population (10^6^ cells) was inoculated on Arabidopsis roots grown under gnotobiotic conditions inside ECO2 boxes in carbon-free silver sand. For each cycle, Arabidopsis seeds were surface sterilized using chlorine gas, germinated on modified Hoagland’s agar medium, and grown for two weeks until transplantation into the ECO2 boxes, containing each two plants (26). Inoculated two-week-old seedlings were then grown for an additional four weeks, after which the root-associated bacteria were collected in 10 mM MgSO_4_ and subjected to fluorescence-based cell counting by flow cytometry yielding on average 10^7^ cells/root. 10^6^ cells of the evolved bacterial populations were then transferred to new plants and this cycle was repeated eight times. After each cycle, a small fraction of each population was plated on general-purpose, nonselective medium, 3 g/l tryptic soy agar (TSA), to assess for contaminations and to verify that all colonies carried the *GFP* marker gene, as observed under UV light, on the one hand, and to select individual isolates for phenotypic characterization on the other hand.

We previously picked sixteen random isolates from each of five experimental lines at cycles 2, 4 and 6 respectively plus sixteen isolates from the ancestor population, yielding a total set of 256 isolates (26). These 256 isolates were subsequently grouped into five distinct phenotypes based on their performance on a variety of bacterial life-history traits (26). In the current study we selected six of these sixteen isolates per line, and per cycle, for genome sequence analysis. Isolates were selected to represent the breath of phenotypic diversity observed previously (26). Together with six isolates from the ancestor population we set out to obtain genome sequences of 96 isolates in total using the NextSeq-500 Illumina platform (2 x 75 bp paired-end). Sequencing of two isolates from line 4, cycle 4, however, failed and thus a final set of 94 genomes were retrieved. We then used the snippy pipeline (https://github.com/tseemann/snippy), integrating reference genome-based mapping of the Illumina reads by BWA-MEM, variant calling by SAMtools and FreeBayes, and variant impact analysis using SnpEff (87), to identify single nucleotide polymorphisms (SNPs) and small indels and deletions (INDELs). Larger INDELs were identified by calculating the breath of coverage of the mapped Illumina reads on the reference genome in a sliding window using bedtools (88). Regions with reduced coverage (<99%) were manually inspected in the Integrative Genome Viewer (IGV). Phylogenetic trees for each line were constructed manually with illustrator based on all detected mutations, the length of the branches representing the number of mutations. The genealogy and frequency of each lineage is shown in the Muller plots that are prepared with R package ‘*ggmuller’*.

### Bacterial life-history traits

For the 94 sequenced isolates, a variety of bacterial life-history traits reflecting various aspects of bacterial physiological processes, were measured previously as part of all 256 isolates initially collected (26). Briefly, we monitored optical density (OD) at a wavelength of 600 nm to estimate the bacterial yield after 72 hours of growth under different growth conditions in 96-well microplates. We measured bacterial growth yield and resistance to various stresses, including acidic (pH = 5) and alkaline (pH = 9) conditions, oxidative stress in 0.0025% H_2_O_2_, water potential stress (15% polyethylene glycol (PEG)-6000)), and salt stress (2% NaCl), and resistance to the antibiotics streptomycin (1 µg · ml^−1^), tetracycline (1 µg · ml^−1^), and penicillin (5 µg · ml^−1^). Bacterial carbon source utilization was quantified as growth yield in modified Ornston and Stanier (OS) minimal medium (89) supplemented with single carbon sources that have been reported to be abundant in Arabidopsis root exudates (Chaparro et al 2013). These included the following carbon sources; alanine, arabinose, butyrolactam, fructose, galactose, glucose, glycerol, glycine, lactic acid, putrescine, serine, succinic acid, threonine, and valine that were added to a final concentration of 0.5 g · l^−1^. In addition, we measured bacterial auxin (indole-3-acetic acid or IAA) production with a colorimetric test (90), iron-chelating ability using a modified chrome azurol S (CAS) assay (91), proteolytic activity by the adapted assay from Smeltzer et al. (1993), tryptophan side chain oxidase activity using a colorimetric assay (92), and biofilm formation using a modified Chrystal Violet staining assay (93). We measured the OD values reflecting the color intensities at specific wavelengths to quantify these traits. We further assessed bacterial antimicrobial activity by quantifying their effect on growth of the fungi *Verticillium dahliae* and *Fusarium oxysporum,* and the bacterium *Ralstonia solanacearum*.

### Motility assays

Motility assays were undertaken in round petri dish plates containing Cook’s Cytophaga medium (CC medium) (0.3% agar for swimming, 0.5% agar for swarming) (94), using typical swimming and swarming assays as described by Déziel et al (95). All tested strains were grown on King’s B medium agar plates for 24 hours before inoculation. Swim and swarm plates were inoculated with the tested strains with a sterile toothpick. For swimming plates, the inoculum was introduced by gently piercing the agar such that the motility within the semisolid agar could be evaluated. For swarming plates, the inoculum was introduced on the agar surface enabling visualization of motility across the agar surface. Both swimming and swarming plates were imaged after 18 hours incubation at 21 °C with right-side-up. The radii of swimming and swarming motility were determined from the photographs by ImageJ, examining the circular turbid zone inside the agar for swimming and the circular zone on the agar surface for swarming.

### Hierarchical and Model-based clustering of bacterial traits

#### Hierarchical clustering

A heatmap to illustrate the association patterns of bacterial genotypes and their measured traits was constructed in R using the *ggplot2* package. Isolates without mutations or only with synonymous mutations were excluded from the association analysis. Hierarchical clustering was performed using the Ward.D2 method, that is based on the squared dissimilarities of the set of objects being clustered (96).

#### Model-based clustering

We applied a model-based clustering method to reveal the best fitting structures of trait covariance patterns. For example, some traits might be either directly or indirectly co-regulated by the same gene, which is expected for global regulators particularly which can co-regulate thousands of genes. We used the *mclust* package in R to run the model simulation (97). This method assumes that the input data are distributed as a mixture of two or more clusters. The advantage of the model-based clustering method is that it avoids heuristic assumptions and uses a soft assignment that every data point has a possibility of falling to each cluster, which facilitates the best clustering solution. The so-called Bayesian Information Criterion (BIC) was used to select the best model. A large BIC score indicates a better fit of the model.

This result is in line with the outcome of hierarchical clustering with Adjusted Rand Index (ARI) set as 1, and *k* set as 4 in “ward.D2” as indicated in Figure 5a. ARI is usually used to evaluate the match degree of a given clustering solution comparing to the model-based clustering result, with 0 reflecting a random partition and 1 the boundary of accuracy of a certain clustering solution (97).

### Genotype-phenotype association analysis

Bacterial traits within each model-predicted cluster have similar data distribution patterns and covaried together by the definition of the clustering method. Thus, we applied a linear regression-based method, *i.e.*, principal component analysis (PCA), to reduce the dimensionality of data and generate a proxy for each model predicted cluster. These proxies were later used as the x-axis values in Figure 5b-e. We applied the package *ggbiplot* in R to generate the PCA plots and PC1 index from the normalized datasets. The proxies were normalized for further analysis.

To examine the accumulative effects of each mutation on bacterial phenotype, ANOVA was used to compare cluster proxies of evolved genotypes with their direct ancestors. Only genotypes identified more than once (*n*>=2) were included in this analysis.

### Relative quantification of mutant frequency using HRM profile analysis

We used High-Resolution Melting (HRM) profile analysis with integrated LunaProbes to quantify the ratio of mutant to wild type genotypes (98–100). The probes and primers used in this study are listed in Table S2. Primers were designed using Primer3. Probes were designed with the single nucleotide polymorphism (SNP) located in the middle of the sequence, and the 3’ end was blocked by carbon spacer C3. The primer asymmetry was set to 2:1 (excess primer: limiting primer) in all cases. Pre-PCR was performed in a 10-μl reaction system, with 0.25 μM excess primer, 0.125 μM limiting primer, 0.25 μM probe, 0.5 μl bacterial sample culture (100-fold diluted saved sample, OD_600_ is about 0.01), 1X LightScanner Master Mix (BioFire Defense). DMSO with the final concentration 5% was supplemented in all reactions to ensure the targeted melting domains are within the detection limit of the LightScanner (Idaho Technology Inc.). Finally, MQ water was used to supplement up to 10 μl. A 96-well black microtiter plate with white wells was used to minimize background fluorescence. Before amplification, 25 μl mineral oil was loaded in each well to prevent evaporation, and the plate was covered with a foil seal to prevent the degradation of fluorescent molecules. Amplification was initiated by a holding at 95 °C for 3 min, followed by 55 cycles of denaturation at 95 °C for 30 s, annealing at 60 °C for 30 s and extension at 72 °C for 30 s and then kept at 72 °C for 10 min. After amplification, samples were heated in a ThermalCycler (Bio-Rad) shortly to 95 °C for 30 s to denature all double-stranded structures followed by a rapid cooling to 25 °C for 30 s to facilitate successful hybridization between probes and the target strands. The plate was then transferred to a LightScanner (Idaho Technology Inc.). Melting profiles of each well were collected by monitoring the continuous loss of fluorescence with a steady increase of the temperature from 35 °C to 97 °C with a ramp rate of 0.1 °C /s. The relative quantification was based on the negative first derivative plots using software MATLAB. The areas of probe-target duplexes melting peaks were auto-calculated by ‘AutoFit Peaks I Residuals’ function in software PeakFit (SeaSolve Software Inc.). The mutant frequency *X* was calculated using the formula shown below:

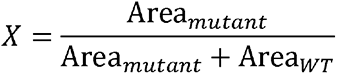

To validate the HRM method, standard curves were generated by measuring mixed samples with known proportions of mutant templates: 0%, 10%, 20%, 30%, 40%, 50%, 60%, 70%, 80%, 90% and 100%. Measurements for each sample were done in triplicate. Linear regression formula of each mutant between actual frequencies and measured frequencies were shown in Figure S1. The high R^2^ values, and nearly equal to 1 slope values of these equations, confirmed that the HRM method can accurately detect mutants’ frequency in a mixed population.

### Data availability

The *P. protegens* CHA0-GFP reference genome is deposited on GenBank: RCSR00000000.1. Raw sequencing data used in this study are deposited at the NCBI database under BioProject PRJNA473919. A conversion table for the CHA0-GFP to CHA0 gene annotations including recent NCBI accession codes is available here: https://doi.org/10.6084/m9.figshare.13295828.v2.

## Acknowledgments

This work was supported by China Scholarship Council fellowships (to E.L. and H.Z.), a postdoctoral fellowship of the Research Foundation Flanders (FWO 12B8116RN) (to R.D.J.), and a European Research Council Advanced Grant 269072 (to C.M.J.P.).

## Author contributions

E.L., H.Z., A.J., C.M.J.P., P.A.H.M.B., and R.D.J. designed experiments; E.L., H.Z., and H.J. performed experiments; E.L., H.Z., and R.D.J. analyzed data. All authors discussed the results and contributed to the final manuscript.

## Supplementary Figure Legends

**Figure S1.** Standard curves of measured mutant versus ancestor proportion as a function of the actual proportion, using series of mixed samples with known proportions (0%, 10%, 20%, 30%, 40%, 50%, 60%, 70%, 80%, 90% and 100% of mutant frequency). Relative densities of mutants *gacA*^D49Y^, *gacA*^D54Y^, and *gacS*^G27D^ were measured by PCR-based high-resolution melting (HRM) analysis. Measurements for each sample were performed in triplicate. In each plot, the black dots represent the measurements, the blue line the fit which was generated based on linear regression modelling.

**Figure S2.** Frequency trajectories of *gacA*^D49Y^, *gacA*^D54Y^ and *gacS*^G27D^ mutants during long-term rhizosphere adaption. The X-axis represents the plant-to-plant transferring cycle of the bacterial population. Mutant frequency was determined by quantifying the ratio of mutant allele relative to wild-type allele, using PCR-based high-resolution melting (HRM) analysis. The data shown are the mean of two technical replicates, and error bars represent the standard deviation of the mean.

**Figure S3.** Representative photographs of swimming (A) and swarming (B) motility assays. Photographs illustrate the experimental layout of motility assays in which every plate is inoculated with the ancestral strain alongside various sequential mutations clustered per experimental line. Genotypes are indicated in each respective corner. All plates were incubated with the right side up at 21 °C for 18 h and then photographed.

**Figure S4.** Selection of the best clustering model using Bayesian Information Criterion (BIC) using the *mclust* R package (97). A large BIC score indicates strong evidence for the corresponding model. The VEI model with 4 components best fits our data.

**Figure S5.** Coordinate axes transformation of each model predicted cluster using principal component analysis (PCA). **a.** PCA plot of social trait cluster (cluster 1); **b.** PCA plot of carbon utilization trait cluster (cluster 2); **c.** PCA plot of biocontrol trait cluster (cluster 3); d. PCA plot of stress resistance trait cluster (cluster 4). Each dot represents one isolate.

## Supplementary Table Legends

**Table S1.** Genotypes of whole-genome sequenced CHA0 isolates.

**Table S2.** Primers and probes used for high-resolution melting (HRM) analysis. For the two gacA mutants the same set of primers was used. Underlined bases indicate the position of the single nucleotide point (SNP) mutations within the probe sequences. ΔTm (°C) indicates the melting temperature difference between WT-probe duplex and mutant-probe duplex.

## Notes

### Competing Interest Statement

The authors have declared no competing interest.

### Summary of Updates

For this revision, in addition to textual improvements, an extended analysis with regards to bacterial motility behavior is included, and we added Muller plots to Figure 3 better illustrating the dynamic of mutations over time.

