## Supplementary figures and images for "Experimental evolution-driven identification of Arabidopsis rhizosphere competence genes in *Pseudomonas protegens*"

### Supplemental Figure 1

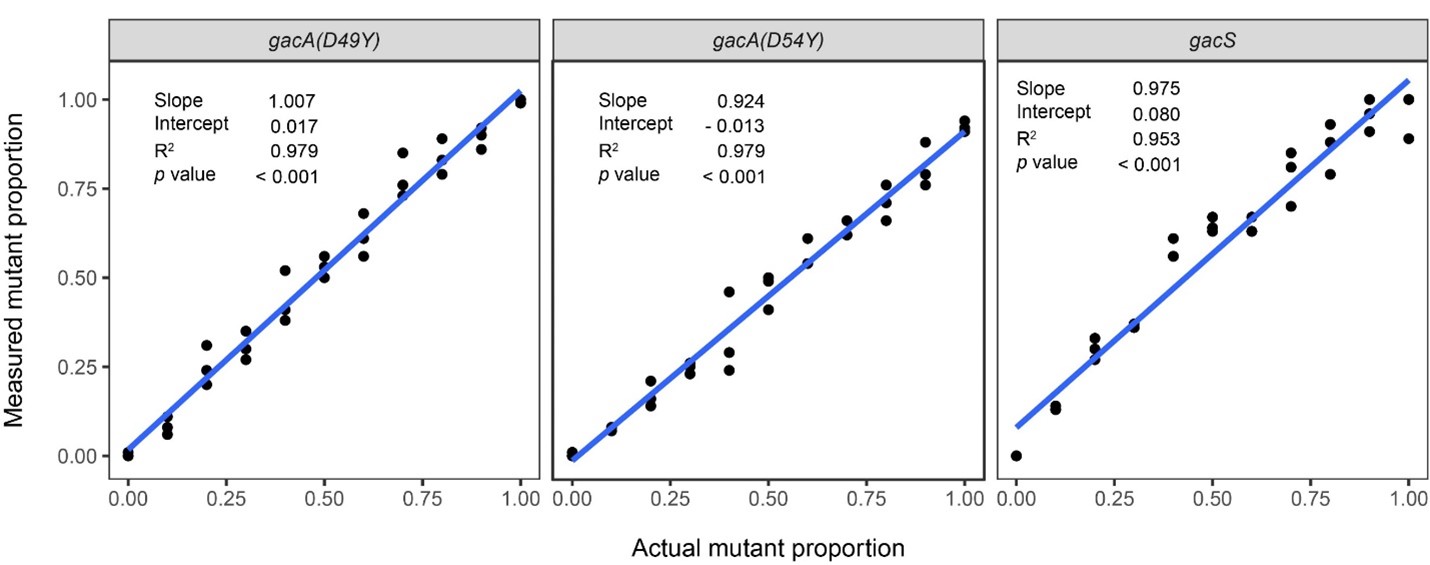

### Supplemental Figure 2

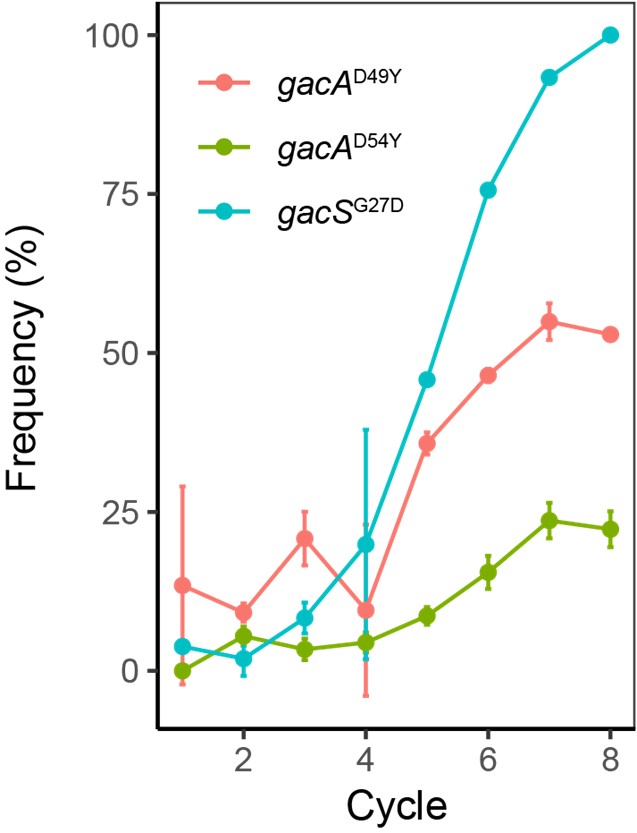

### Supplemental Figure 3

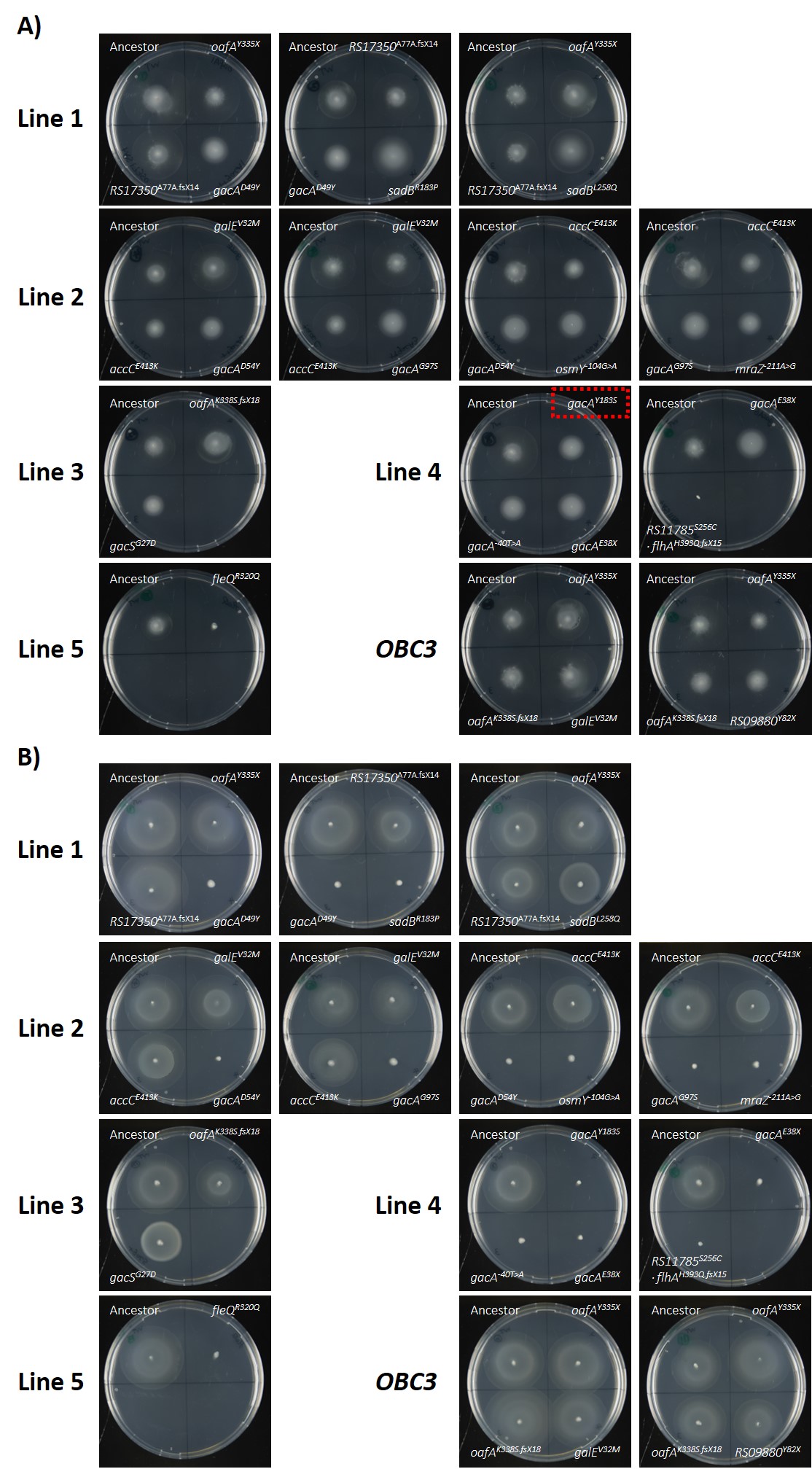

### Supplemental Figure 4

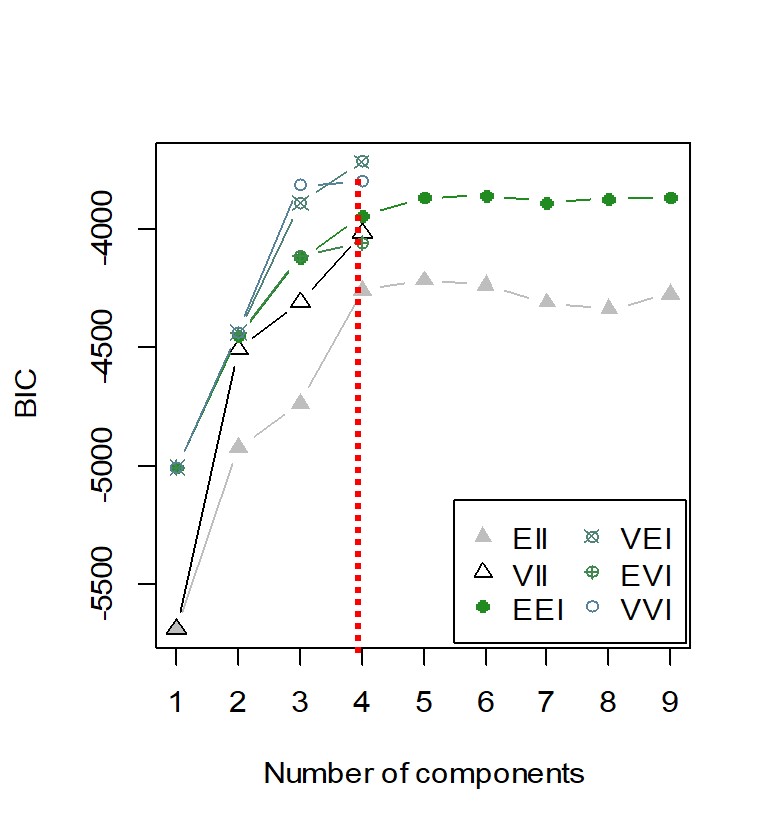

### Supplemental Figure 5

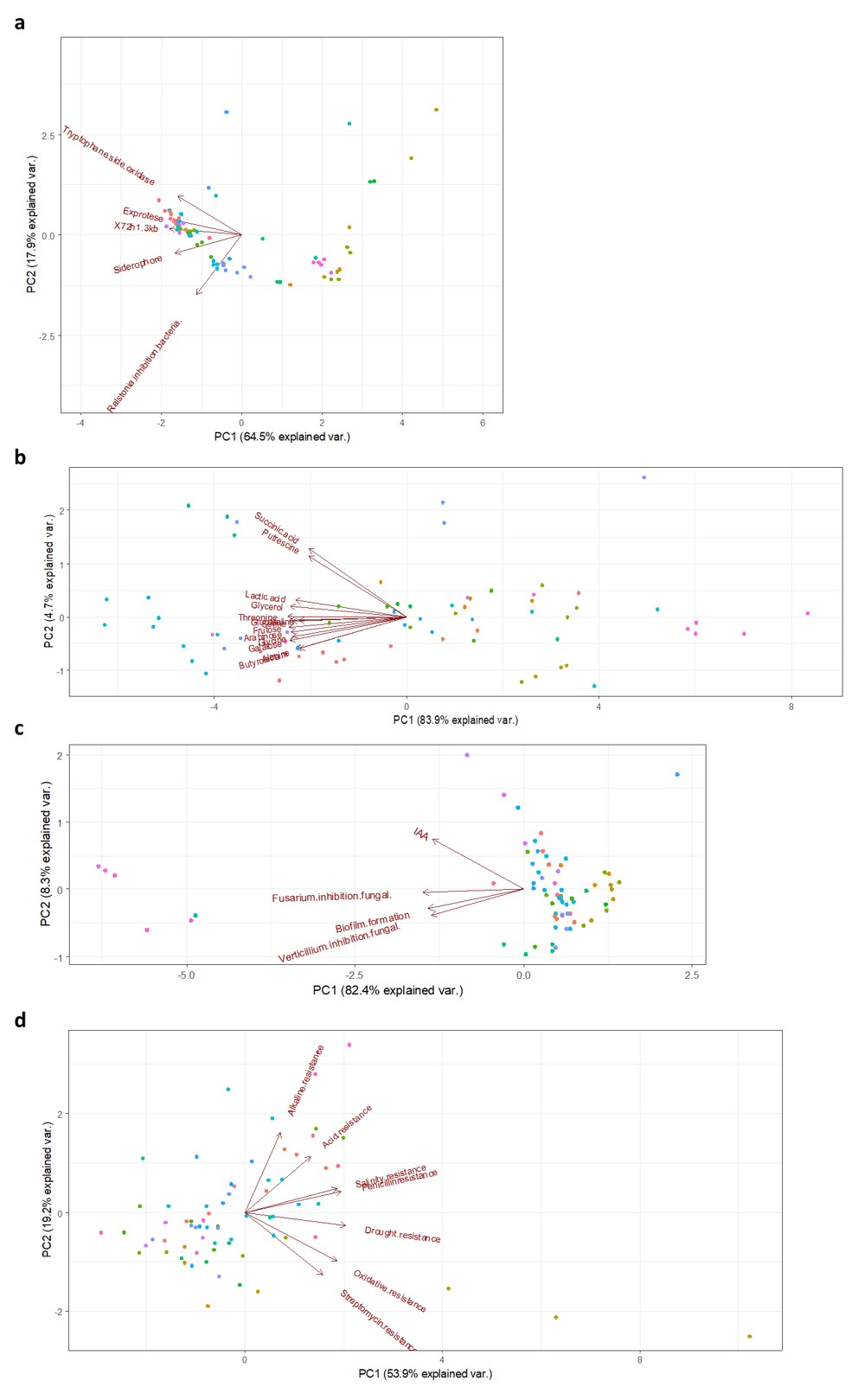
