## Supplemental Table 1 for "Experimental evolution-driven identification of Arabidopsis rhizosphere competence genes in *Pseudomonas protegens*"

**Table S1** Genotypes of whole-genome sequenced CHA0 isolates.

| **Sample ID** | **Line** | **Cycle** | **Genotype** | **NCBI SRR IDENTIFIER** |
| --- | --- | --- | --- | --- |
| 61 | 1 | 2 | *WT* | SRR13155042 |
| 169 | 1 | 2 | *WT* | SRR13155041 |
| 13 | 1 | 2 | *oafA*^Y335X^ | SRR13155030 |
| 121 | 1 | 2 | *oafA*^Y335X^ | SRR13155019 |
| 133 | 1 | 2 | *oafA*^Y335X^ *∙ RS17350*^A77A.fsX14^ | SRR8053478 |
| 157 | 1 | 2 | *oafA*^Y335X^ *∙ RS17350*^A77A.fsX14^ | SRR13155008 |
| 158 | 1 | 4 | *oafA*^Y335X^ *∙ RS17350*^A77A.fsX14^ | SRR13154997 |
| 182 | 1 | 4 | *oafA*^Y335X^ *∙ RS17350*^A77A.fsX14^ | SRR13154986 |
| 14 | 1 | 4 | *oafA*^Y335X^ *∙ RS17350*^A77A.fsX14^ *∙ gacA*^D49Y^ | SRR13154981 |
| 122 | 1 | 4 | *oafA*^Y335X^ *∙ RS17350*^A77A.fsX14^ *∙ gacA*^D49Y^ | SRR13154980 |
| 86 | 1 | 4 | *oafA*^Y335X^ *∙ RS17350*^A77A.fsX14^ *∙ wbpM*^G79R^ | SRR8053477 |
| 2 | 1 | 4 | *oafA*^Y335X^ *∙ RS17350*^A77A.fsX14^ *∙ RS21275*^R78L^ | SRR13154979 |
| 195 | 1 | 6 | *oafA*^Y335X^ *∙ RS17350*^A77A.fsX14^ *∙ gacA*^D49Y^ | SRR13155040 |
| 242 | 1 | 6 | *oafA*^Y335X^ *∙ RS17350*^A77A.fsX14^ *∙ gacA*^D49Y^ | SRR8053476 |
| 278 | 1 | 6 | *oafA*^Y335X^ *∙ RS17350*^A77A.fsX14^ *∙ gacA*^D49Y^ | SRR13155039 |
| 255 | 1 | 6 | *oafA*^Y335X^ *∙ RS17350*^A77A.fsX14^ *∙ gacA*^D49Y^ *∙ sadB*^R183P^ | SRR13155038 |
| 206 | 1 | 6 | *oafA*^Y335X^ *∙ RS17350*^A77A.fsX14^ *∙ sadB*^L258Q^ | SRR13155037 |
| 207 | 1 | 6 | *oafA*^Y335X^ *∙ RS17350*^A77A.fsX14^ *∙ sadB*^L258Q^ *∙ pvdS*^-33T>C^ | SRR13155036 |
| 87 | 2 | 2 | *WT* | SRR13155035 |
| 39 | 2 | 2 | *gacA*^Y183S^ | SRR13155034 |
| 183 | 2 | 2 | *gacA*^Y183S^ | SRR13155033 |
| 123 | 2 | 2 | *galE*^V32M^ | SRR13155032 |
| 135 | 2 | 2 | *galE*^V32M^ | SRR13155031 |
| 63 | 2 | 2 | *tssM*^1336C>T^ | SRR13155029 |
| 172 | 2 | 4 | *gacA*^Y183S^ | SRR8053471 |
| 52 | 2 | 4 | *galE*^V32M^ | SRR8053473 |
| 16 | 2 | 4 | *galE*^V32M^ *∙ accC*^E413K^ | SRR8053475 |
| 28 | 2 | 4 | *galE*^V32M^ *∙ accC*^E413K^ | SRR8053474 |
| 100 | 2 | 4 | *galE*^V32M^ *∙ accC*^E413K^ | SRR13155028 |
| 160 | 2 | 4 | *galE*^V32M^ *∙ accC*^E413K^ *∙ yvaQ2*^-9G>T^ | SRR8053472 |
| 245 | 2 | 6 | *galE*^V32M^ *∙ accC*^E413K^ | SRR13155027 |
| 220 | 2 | 6 | *galE*^V32M^ *∙ accC*^E413K^ *∙ gacA*^D54Y^ | SRR8053470 |
| 197 | 2 | 6 | *galE*^V32M^ *∙ accC*^E413K^ *∙ gacA*^D54Y^ *∙ osmY*^-104G>A^ | SRR13155026 |
| 221 | 2 | 6 | *galE*^V32M^ *∙ accC*^E413K^ *∙ gacA*^G97S^ | SRR13155025 |
| 196 | 2 | 6 | *galE*^V32M^ *∙ accC*^E413K^ *∙ gacA*^G97S^ *∙ mraZ*^-211A>G^ | SRR13155024 |
| 268 | 2 | 6 | *galE*^V32M^ *∙ accC*^E413K^ *∙ gacA*^G97S^ *∙ mraZ*^-211A>G^ | SRR8053469 |
| 29 | 3 | 2 | *WT* | SRR13155023 |
| 65 | 3 | 2 | *WT* | SRR8053484 |
| 53 | 3 | 2 | *WT* | SRR13155022 |
| 77 | 3 | 2 | *WT* | SRR13155021 |
| 161 | 3 | 2 | *WT* | SRR13155020 |
| 137 | 3 | 2 | *oafA*^K338S.fsX18^ | SRR13155018 |
| 90 | 3 | 4 | *nlpD*^Q197P^ | SRR8053486 |
| 18 | 3 | 4 | *oafA*^K338S.fsX18^ | SRR13155017 |
| 66 | 3 | 4 | *oafA*^K338S.fsX18^ | SRR8053483 |
| 150 | 3 | 4 | *oafA*^K338S.fsX18^ | SRR13155016 |
| 186 | 3 | 4 | *oafA*^K338S.fsX18^ | SRR13155015 |
| 30 | 3 | 4 | *oafA*^K338S.fsX18^ *∙ gacS*^G27D^ | SRR13155014 |
| 235 | 3 | 6 | *oafA*^K338S.fsX18^ | SRR13155013 |
| 270 | 3 | 6 | *oafA*^K338S.fsX18^ | SRR13155012 |
| 199 | 3 | 6 | *oafA*^K338S.fsX18^ *∙ gacS*^G27D^ | SRR13155011 |
| 222 | 3 | 6 | *oafA*^K338S.fsX18^ *∙ gacS*^G27D^ | SRR8053485 |
| 234 | 3 | 6 | *oafA*^K338S.fsX18^ *∙ gacS*^G27D^ | SRR13155010 |
| 198 | 3 | 6 | *oafA*^K338S.fsX18^ *∙ gacS*^G27D^ | SRR13155009 |
| 57 | 4 | 2 | *WT* | SRR13155007 |
| 105 | 4 | 2 | *WT* | SRR13155006 |
| 81 | 4 | 2 | *WT* | SRR13155005 |
| 129 | 4 | 2 | *∆argT5* | SRR13155004 |
| 141 | 4 | 2 | *∆argT5* | SRR13155003 |
| 21 | 4 | 2 | *hult*^786C>T^ | SRR8053480 |
| 80 | 4 | 4 | *WT* | SRR8053482 |
| 152 | 4 | 4 | *WT* | SRR13155002 |
| 188 | 4 | 4 | *gacA*^-40T>A^ | SRR8053481 |
| 68 | 4 | 4 | *gacA*^-40T>A^ *∙ RS11820*^33C>T^ | SRR8053479 |
| 164 | 4 | 4 | *gacA*^E38X^ | SRR13155001 |
| 20 | 4 | 4 | *gacA*^E38X^ *∙*  *nudL*^288C>T^ | SRR13155000 |
| 225 | 4 | 6 | *gacA*^-40T>A^ *∙ rpsH*^-90C>G^ | SRR13154999 |
| 212 | 4 | 6 | *gacA*^E38X^ *∙ RS11785*^S256C^ *∙ flhA*^H393Q.fsX15^ | SRR13154998 |
| 237 | 4 | 6 | *gacA*^E38X^ *∙ RS11785*^S256C^ *∙ flhA*^H393Q.fsX15^ | SRR13154996 |
| 260 | 4 | 6 | *gacA*^E38X^ *∙ RS11785*^S256C^ *∙ flhA*^H393Q.fsX15^ | SRR8053488 |
| 11 | 6 | 2 | *WT* | SRR13154995 |
| 71 | 6 | 2 | *WT* | SRR13154994 |
| 191 | 6 | 2 | *WT* | SRR13154993 |
| 35 | 6 | 2 | *WT* | SRR13154992 |
| 83 | 6 | 2 | *WT* | SRR13154991 |
| 143 | 6 | 2 | *RS18525*^V85G.fsX40^ | SRR13154990 |
| 70 | 6 | 4 | *WT* | SRR8053461 |
| 106 | 6 | 4 | *WT* | SRR13154989 |
| 142 | 6 | 4 | *WT* | SRR13154988 |
| 58 | 6 | 4 | *WT* | SRR8053487 |
| 190 | 6 | 4 | *WT* | SRR13154987 |
| 34 | 6 | 4 | *rpoS*^Q65X^ | SRR13154985 |
| 251 | 6 | 6 | *fleQ*^R320Q^ | SRR8053463 |
| 262 | 6 | 6 | *rpoS*^Q65X^ | SRR8053464 |
| 263 | 6 | 6 | *rpoS*^Q65X^ | SRR8053465 |
| 203 | 6 | 6 | *rpoS*^Q65X^ *∙ tetR*^Y127X^ | SRR13154984 |
| 239 | 6 | 6 | *rpoS*^Q65X^ *∙ tetR*^Y127X^ | SRR8053462 |
| 226 | 6 | 6 | *RS09880*^Y82X^ *∙ RS12070*^1389C>G^ | SRR13154983 |
| 7 | ancestor | 0 | *WT* | SRR8053466 |
| 31 | ancestor | 0 | *WT* | SRR8053467 |
| 79 | ancestor | 0 | *WT* | SRR8053468 |
| 127 | ancestor | 0 | *WT* | SRR8053459 |
| 163 | ancestor | 0 | *WT* | SRR8053460 |
| 187 | ancestor | 0 | *WT* | SRR13154982 |

Note: X, represents a stop codon (at its relative position in case of a shifted frame); fs, frame shift; del, deletion; DNA sequence change positions are relative to the cDNA.
