## Supplemental Table 2 for "Experimental evolution-driven identification of Arabidopsis rhizosphere competence genes in *Pseudomonas protegens*"

**Table S2** Primers and probes used for high-resolution melting (HRM) analysis. For the two *gacA* mutants the same set of primers was used. Underlined bases indicate the position of the single nucleotide point (SNP) mutations within the probe sequences. ∆Tm (℃) indicates the melting temperature difference between WT-probe duplex and mutant-probe duplex.

| **Target gene** | **Strain ID** | **SNP locus** | **Forward primer (excess)** | **Reverse primer (limiting)** | **Amplicon size** | **Probe sequence** | **Probe length** | **Target strand** | **Perfect match/ mismatch** | **_∆_Tm (℃)** |
| --- | --- | --- | --- | --- | --- | --- | --- | --- | --- | --- |
| *gacA*^D49Y^ | 242 | 145G>T | 5'-ATCGATGGCCTGCAAGTAGT-3’ | 5'-CGGGTAGGAAAGGGATCTTC-3’ | 206 bp | 5’-CATCAGGACCACATCGGGCTTCAGCTCCCG-/C3/3’ | 30nt | WT | G::C / T::C | 5.31 |
| *gacA*^D54Y^ | 220 | 160G>T |  |  |  | 5’-TGGCATCTTGACGTCCATCAGGACCACATC-/C3/3’ | 30nt | WT | G::C / T::C | 4.92 |
| *gacS*^G27D^ | 222 | 80G>A | 5’-GCGTACTGTTGCTGACCTTG-3’ | 5’-AGCATCTGGGTGTTGTGGTT-3’ | 178bp | 5’-AGGTGAAGTAGCCGCCCAGCACCAAAGCCA-/C3/3’ | 30nt | WT | G::C / A::C | 4.83 |
